# Cell Atlas of the Human Fovea and Peripheral Retina

**DOI:** 10.1101/2020.02.11.943779

**Authors:** Wenjun Yan, Yi-Rong Peng, Tavé van Zyl, Aviv Regev, Karthik Shekhar, Dejan Juric, Joshua R Sanes

## Abstract

Most irreversible blindness results from retinal disease. To advance our understanding of the etiology of blinding diseases, we used single-cell RNA-sequencing (scRNA-seq) to analyze the transcriptomes of ∼85,000 cells from the fovea and peripheral retina of seven adult human donors. Utilizing computational methods, we identified 58 cell types within 6 classes: photoreceptor, horizontal, bipolar, amacrine, retinal ganglion and non-neuronal cells. Nearly all types are shared between the two retinal regions, but there are notable differences in gene expression and proportions between foveal and peripheral cohorts of shared types. We then used the human retinal atlas to map expression of 636 genes implicated as causes of or risk factors for blinding diseases. Many are expressed in striking cell class-, type-, or region-specific patterns. Finally, we compared gene expression signatures of cell types between human and the cynomolgus macaque monkey, *Macaca fascicularis*. We show that over 90% of human types correspond transcriptomically to those previously identified in macaque, and that expression of disease-related genes is largely conserved between the two species. These results validate the use of the macaque for modeling blinding disease, and provide a foundation for investigating molecular mechanisms underlying visual processing.

## INTRODUCTION

The three leading causes of irreversible blindness can be classified as neurodegenerative retinal diseases: age-related macular degeneration, glaucoma and diabetic retinopathy; photoreceptors are lost in age-related macular degeneration and diabetic retinopathy, and retinal ganglion cells (RGCs) are lost in glaucoma (1–3). These three groups of diseases affect over 300 million people world-wide, greatly outnumbering those affected by other neurodegenerative diseases such as Alzheimer’s and Parkinson’s diseases. Genetic contributors have been discovered for all of these retinal diseases, largely through genome-wide association studies (GWAS) (4–6). In few cases, however, do we understand the role of the implicated gene in disease pathogenesis.

One main obstacle to gaining such understanding is lack of knowledge about where the implicated genes are expressed; lacking such information, it is difficult to determine mechanisms by which it affects visual function. Another is that substantial differences in structure and gene expression between human and rodent retina have made it difficult to study these genes in animal models. For example, among mammals, only primates have a fovea, the small central region responsible for high acuity vision as well as most chromatic vision – and the region selectively affected in macular degeneration, diabetic macular edema and hereditary maculopathies (7, 8). As a first step toward addressing these issues, we recently used high throughput single cell RNA-seq (scRNA-seq) to generate a retinal cell atlas from cynomolgus macaque (*Macaca fascicularis*), a non-human primate that is closely related to humans and frequently used in preclinical ophthalmological studies (9–11). We separately profiled peripheral retina and the fovea (Fig.1a). In each region, we characterized the six classes of retinal cells – photoreceptors, horizontal cells, bipolar cells, amacrine cells, RGCs and Müller glial cells (Fig. 1b) and found molecular signatures that divided them into a total of 62 (fovea) or 70 (periphery) cell types (12).

**Figure 1.**
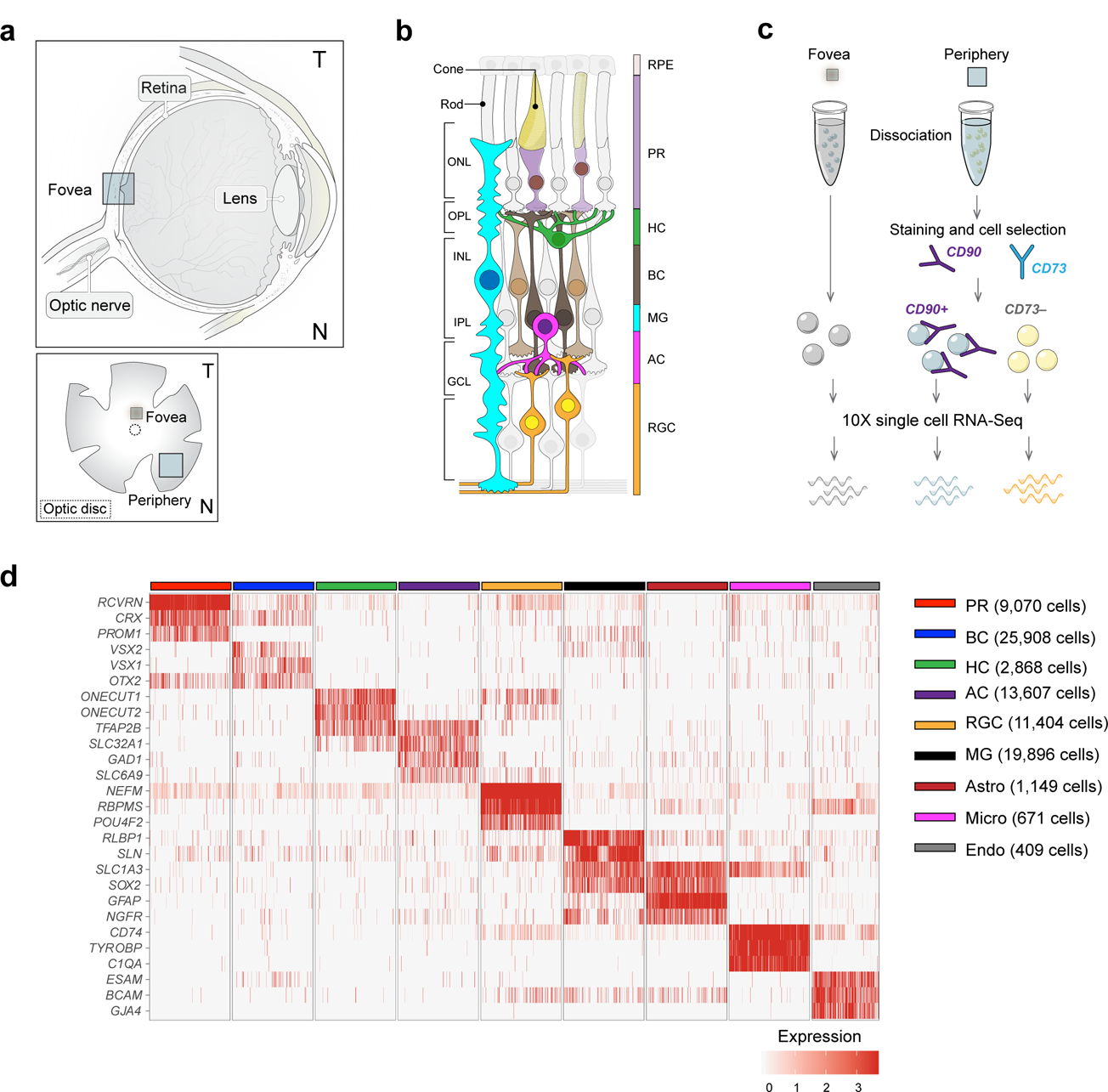
Cell classes in human retina. (a) (Top) Sketch of a human eye showing positions of retina and fovea. (Bottom) Sketch of a flat mounted retina showing foveal and peripheral regions. Circle in center is position at which optic nerve exits the eye. (b) Sketch of a peripheral retinal section showing its major cell classes—photoreceptors (PRs), horizontal cells (HCs), bipolar cells (BCs), amacrine cells (ACs), retinal ganglion cells (RGCs) and Müller glia (MG), outer and inner plexiform (synaptic) layers (OPL and IPL), outer and inner nuclear layers (ONL and INL), and ganglion cell layer (GCL). (c) scRNA-seq workflow. Foveal cells were dissociated from < 1.5 mm-diameter punches and collected without further processing. Peripheral cells were dissociated from all four quadrants of peripheral retinas, and depleted of rods (CD73+) or enriched for RGCs (CD90+) with magnetic columns before processing. (d) Expression patterns of class-specific markers (rows) in individual cells (columns). Cells are grouped by their classes (color bars, top). Plot shows data from a maximum of 500 randomly selected cells per class.

Here, to extend this work, we used the macaque atlas as a foundation to generate a comprehensive cell atlas of the adult human retina. We expected that the similarity of macaque to human would aid in identifying cell types, and this was indeed the case. By analyzing a total of 84,982 single cell transcriptomes, we identified 58 cell types in human fovea and 57 types in peripheral retina, nearly all of which were shared between the two regions. For many of these types, however, we documented substantial regional differences in gene expression and proportions. By comparing human and macaque atlases, we found 1:1 matches for >90% of cell types, supporting the use of *Macaca fascicularis* as a preclinical model. Finally, we mapped the expression of 636 genes implicated in blinding diseases by GWAS studies or as highly penetrant Mendelian mutations underlying a variety of inherited retinal degenerations, each rare but substantial in aggregate. We show that many of the genes queried are selectively expressed in particular retinal cell classes, in particular cell types within a class, or in foveal or peripheral cohorts of shared types. These results provide new insights into mechanisms underlying retinal disease.

## RESULTS

### Cell classes in human retina

To generate a comprehensive cell atlas of human retina, we obtained eight retinas from seven genetically unrelated human donors with no clinical history of ocular disease (Table S1). We dissected fovea (∼1.5 mm diameter centered on the foveal pit, which was visible under a dissecting microscope) and peripheral samples (> 5 mm from the fovea) from whole retina, pooling peripheral pieces from all four quadrants. Foveal samples were dissociated into single cells, which were profiled without further processing using high-throughput droplet sequencing (13). For peripheral samples, in which rod photoreceptors and RGC comprise ∼80% and <2 % of total cells respectively, we depleted rods using magnetic beads conjugated to anti-CD73 or enriched RGCs using anti-CD90-conjugated beads prior to collection (Fig.1c), using protocols established in our study on macaque retina (12). Libraries were prepared from foveal and peripheral samples, and sequenced. Altogether, we obtained 84,982 high-quality transcriptomes, 55,736 from fovea and 29,246 from peripheral retina. The median number of unique transcripts captured per cell was 2,577 and the median number of genes detected was 1,308.

To maximize statistical power, we pooled data from fovea and periphery for initial analysis. Using methods adapted from (12), we divided the cells into 9 groups based on expression of canonical markers, which were common to both retinal regions (Fig. 1d). We identified the five neuronal classes (9,070 photoreceptors, 2,868 horizontal cells, 25,908 bipolar cells, 13,607 amacrine cells and 11,404 RGCs) as well as four types of non-neuronal cells: 19,896 Müller glia, 1,149 astrocytes, 671 microglia and 409 vascular endothelial cells.

### Classification and identification of retinal cell types

We next re-clustered each neuronal class separately to discriminate cell types. We obtained a total of 54 clusters, each corresponding to a putative cell type or possibly a small group of closely related types: 3 photoreceptor, 2 horizontal cell, 12 bipolar cell, 25 amacrine cell, and 12 RGC types. Thus, including the 4 non-neuronal types, we detected a total of 58 cell types in human retina. Of them, 54 contained cells from at least 6 of the 7 donors (Supplemental Fig. 1), indicating that the heterogeneity does not result from individual variations or batch effects. We took advantage of the evolutionary proximity between humans and macaques and utilized previously defined macaque retina cell types (12) to train a multi-class supervised classification algorithm (14). This enabled us to relate most human clusters to macaque types, based on their expression patterns of orthologous genes. Many of the human types were further characterized by assessing their expression of key genes reported previously.

#### Photoreceptors

The two subclasses of photoreceptor cells in vertebrate retinas are rods, specialized for high-sensitivity vision at low light levels, and cones, which mediate chromatic vision. Rods and cones express rhodopsin and cone opsins, respectively. Humans and many old world monkeys, such as macaques, are trichromats, with three cone types, each expressing a single opsin (S-, M- or L-opsin) tuned to short-, medium- or long-wavelengths, respectively. We found three clear photoreceptor clusters: rods, which selectively express rhodopsin; S-cones, which selectively express S-opsin; and M and L cones, which selectively express and M or L-opsin (Fig. 2a-c). The inability to distinguish M from L opsin results from their nearly identical coding sequences (98% nucleotide identity), the presence of multiple copies of the M-opsin gene in some individuals, and the high frequency of recombination between these two closely linked genes (see Peng et al., 2019 for discussion). The three human photoreceptor types mapped to their macaque counterparts with high confidence (Fig. 2b).

**Figure 2.**
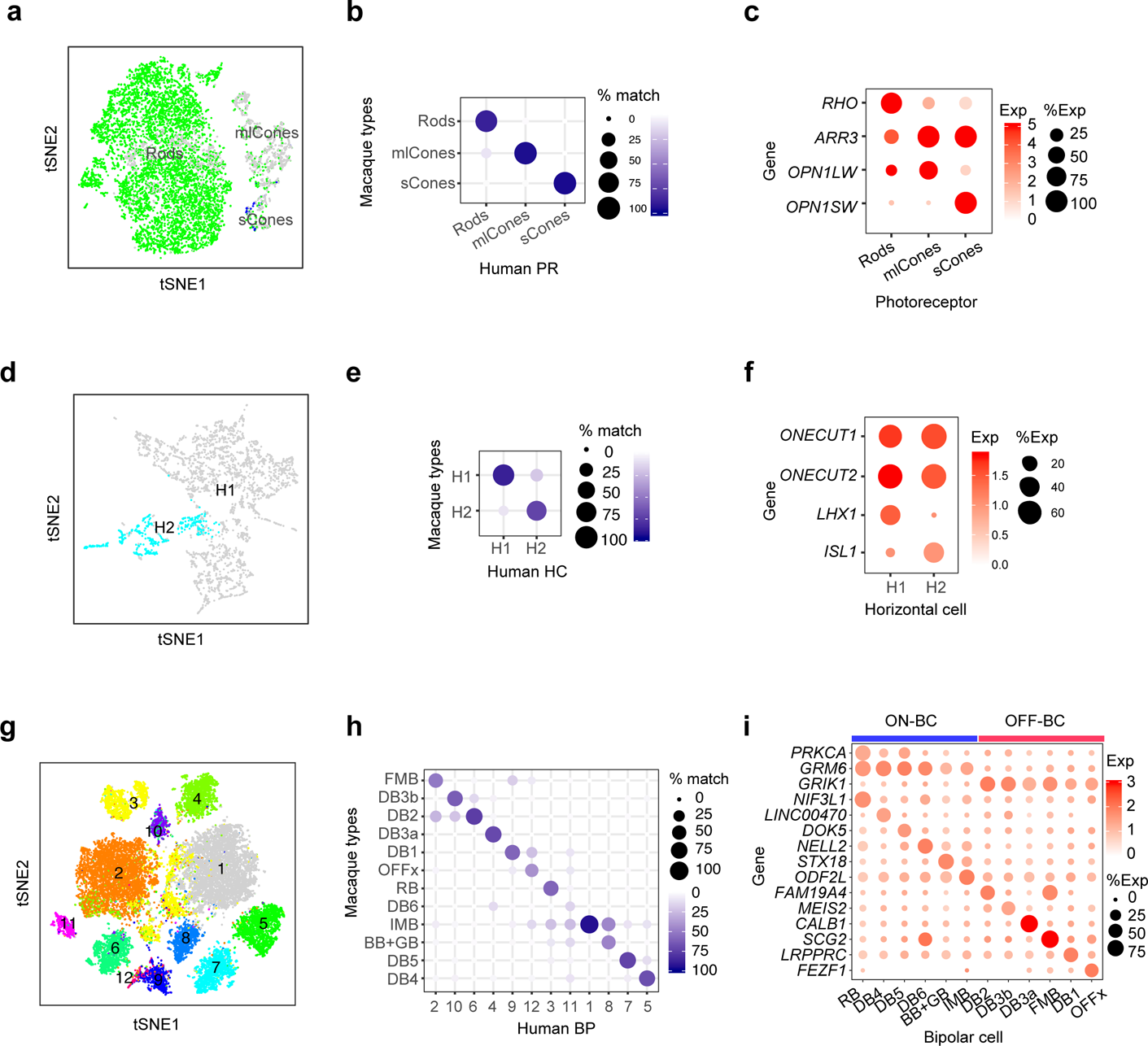
Photoreceptor, horizontal and bipolar cell types. (a-c). Photoreceptors. (a) Cell clusters visualized using t-distributed stochastic neighbor embedding (t-SNE). Dots represent individual cells and are color coded by their cluster assignments (text labels). (b) Transcriptional correspondence between human and macaque cell types summarized as a “confusion matrix.” In this and subsequent confusion matrices, the color and size of each dot reflect the percentage of cells in a given human cluster (columns) mapped to a corresponding macaque type (rows). (c) Dot plots showing expression of select marker genes. In this and subsequent gene expression plots, dot size represents the percentage of cells in a cluster with non-zero expression of a select gene; and color intensity represents the average expression of a gene within expressing cells. (d-f) Horizontal cells. Clusters (d), correspondence to macaque types (e) and expression of key genes (f) shown as in a-c. (g-i) Bipolar cells. Clusters (g), correspondence to macaque types (h) and expression of key genes (i) shown as in a-c.

#### Horizontal cells

Most primates, including macaques, have two horizontal cell types, H1 and H2 (15). Based on morphological criteria, Kolb and colleagues argued for a third horizontal cell type in human retina (16, 17), with H3 differing from H1 in having larger somata and dendritic arbors. We identified two horizontal cell types (Fig. 2d), which corresponded to the macaque H1 and H2 types, respectively (Fig. 2e). Attempts to further subdivide the two types by increasing the sensitivity of the clustering algorithm failed to reveal a third type with a distinguishing molecular signature. Similar to macaque H1 and H2, human H1 and H2 were distinguished from each other by selectively expressing transcription factors *LHX1* and *ISL1*, respectively (Fig. 2f).

#### Bipolar cells

Bipolar cells are divided into three subclasses: ON and OFF cone bipolars, which release neurotransmitter in response to increases and decreases in illumination of cones, respectively; and rod bipolars, which generate ON responses to stimulation of rods (18, 19). In macaques, ON and OFF bipolars are characterized by expression of genes that encode the metabotropic glutamate receptor 6, *GRM6*, and the kainate-type glutamate receptor, *GRIK1*, respectively; rod bipolars are distinguished from cone bipolars by expression of *PKCα*, encoding protein kinase C*α* (12). These expression patterns were conserved in human bipolars, allowing us to divide 12 bipolar clusters into 1 ON rod, 5 ON cone, and 6 OFF cone types (Fig. 2g-i). The counterparts of all 12 macaque types were found in human retina and named based on this correspondence (Fig. 2h). Notably, the provisionally named “OFFx” type, first identified and named in our analysis of macaque retina, was also present in human retina as a distinct cluster (Fig. 2h, i).

#### Amacrine cells

Most amacrine cells are inhibitory neurons utilizing GABA or glycine as neurotransmitters. By assessing the expression of marker genes for GABAergic (glutamate carboxylase, *GAD1* and *GAD2*) and glycinergic (*SLC6A9*, encoding the high affinity glycine transporter GLYT1) amacrines (20), we identified 16 putative GABAergic and 8 putative glycinergic amacrine cell types among a total of 25 types (Fig. 3a, b). One type (C14) expressed none of these three genes at high levels, and might correspond to a non-GABAergic non-Glycinergic (nGnG) type identified in mouse (21; Yan et al., in preparation). One of the glycinergic types (C17) also expressed *GAD2*, raising the possibility that it uses both transmitters. Several known amacrine types were detected based on key marker genes (Fig. 3d), including *SLC17A8* for VG3 amacrine (an excitatory type that co-releases glycine and glutamate), *SLC18A3* for cholinergic starburst amacrines, TH for catecholaminergic CAI/CAII amacrines, and *GJD2* for AII amacrines, which mediate transmission of rod signals to RGCs (22). Those and many other AC types mapped faithfully to macaque types (Fig. 3c). Many AC types also expressed neuropeptides (bold in Fig. 3d), with some predominantly in single types (e.g. *NPW* in C7 and *VIP* in C24, and others expressed by multiple types (e.g. *CARTPT* and *PENK*). In several instances, more than one neuropeptide was detected in the same AC type – for example, thyrotropin-releasing hormone (*TRH*) and Natriuretic Peptide B (*NPPB*) in C9, and Proenkephalin (*PENK*) and Cholecystokinin (*CCK*) in C15. Thus, human amacrines appear to use a variety of neurotransmitters and neuromodulators, as has been demonstrated for amacrines in other mammalian and non-mammalian retinas (W.Y. and J.R.S., in preparation).

**Figure 3.**
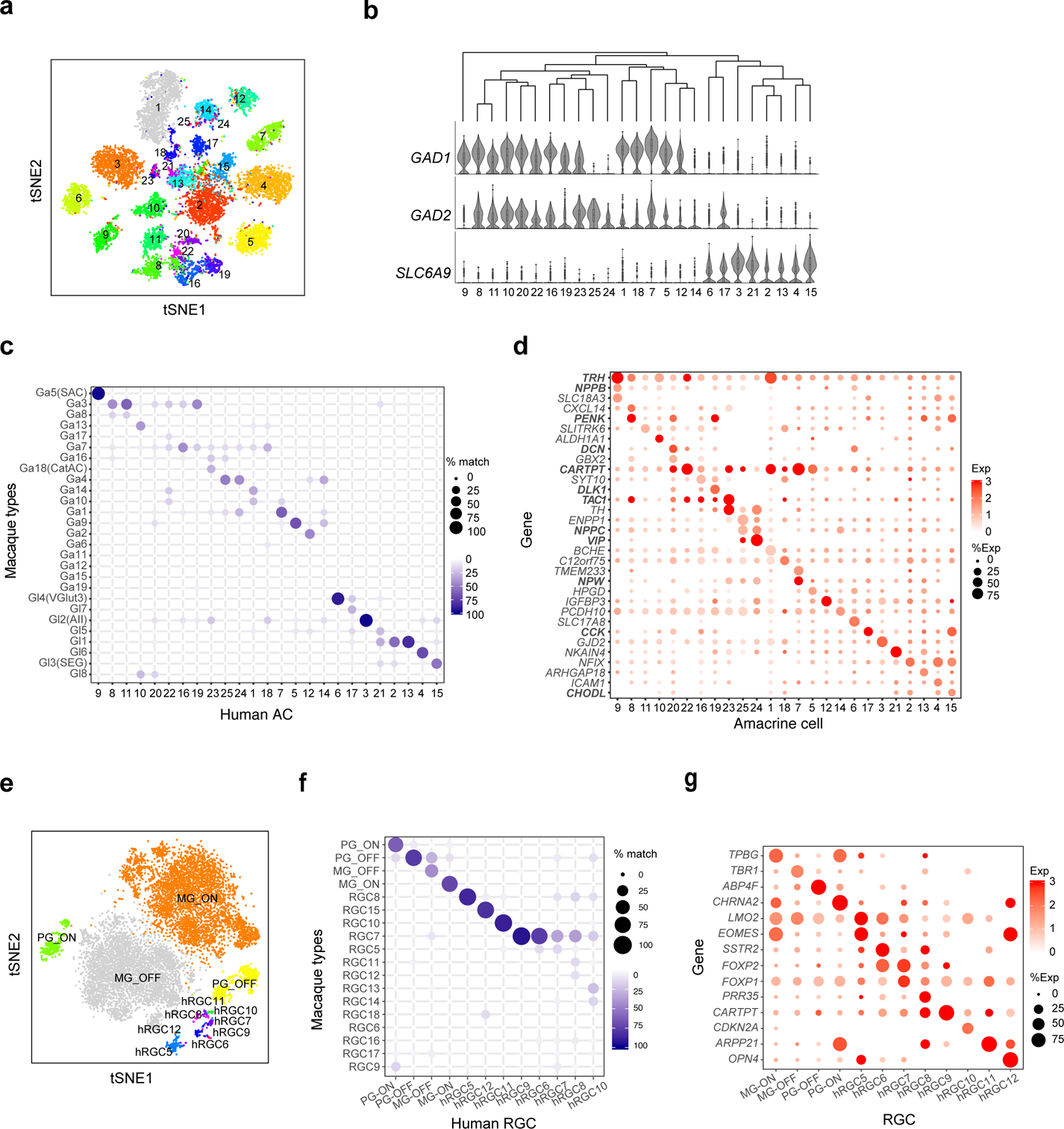
Amacrine and retinal ganglion cell types. (a-d) Amacrine cells. Clusters visualized by tSNE (a), correspondence to macaque types (c) and expression of key genes (d) shown as in Figure 2. Known amacrine types (SAC, VG3-AC, Aii, SEG) are conserved between macaque and human. Genes encoding neuropeptides are shown in bold in d. (b) Top, dendrogram showing transcriptomic relationships among AC clusters; Bottom, violin plots representing the distribution of expression of GABAergic (*GAD1, GAD2*) and Glycinergic (*SLC6A9*) in each AC cluster. (e-g) Retinal ganglion cell clusters visualized by tSNE (e), correspondence to macaque types (f) and expression of key genes (g) shown as in Figure 2.

*Retinal ganglion cells*

The predominant ganglion cell types in primate retina are ON and OFF midget RGCs, together accounting for >80% of RGCs in human (by morphological criteria) and macaque retina (by morphological and molecular criteria) (12, 23). Next most abundant in both species are ON and OFF parasol RGCs, totaling ∼10% of all RGCs. Based on abundance, four RGC clusters appeared likely to correspond to these types (Fig. 3e). Mapping to the macaque atlas confirmed their identities (Fig. 3f). The midget and parasol RGCs comprised 86% (44% ON and 42% OFF) and 10% (4% ON and 6% OFF) of all RGCs in our dataset, respectively.

The remaining 8 clusters ranged in abundance from 0.1% to 1.6% of all RGCs. They included two types that expressed the transcription factor, *FOXP2* (hRGC6 and 7), one of which also expressed *FOXP1* (Fig. 3g); these might be related to mouse FoxP2+FoxP1- and FoxP2+FoxP1+ F-RGCs (24). We also detected two RGC clusters that expressed melanopsin (*OPN4*), the canonical marker of intrinsically photosensitive RGCs (ipRGCs; hRGC5 and hRGC12; Fig. 3g). Recent morphological and physiological studies have demonstrated 2-4 human ipRGC types (25, 26). We speculate that hRGC12, which expressed the highest level of *OPN4* (Fig. S2) corresponds to M1, which expresses highest levels of *OPN4* in mice (27); others could be included in hRGC5 or be too rare to detect.

*Non-neuronal cells*.

Four clusters of non-neuronal cells were identified as Müller glia, astrocytes, microglia, and endothelial cells based on expression of known markers (Fig. 1d). The Müller cell, the intrinsic glial cell of the retina, was the most abundant type among them (Fig. 4a). Astrocytes, which are largely confined to the ganglion cell and nerve fiber layers, were transcriptomically similar to Müller glia (Fig. 1d), but the two types were readily distinguished by selective expression of multiple genes (Fig. 4b).

**Figure 4.**
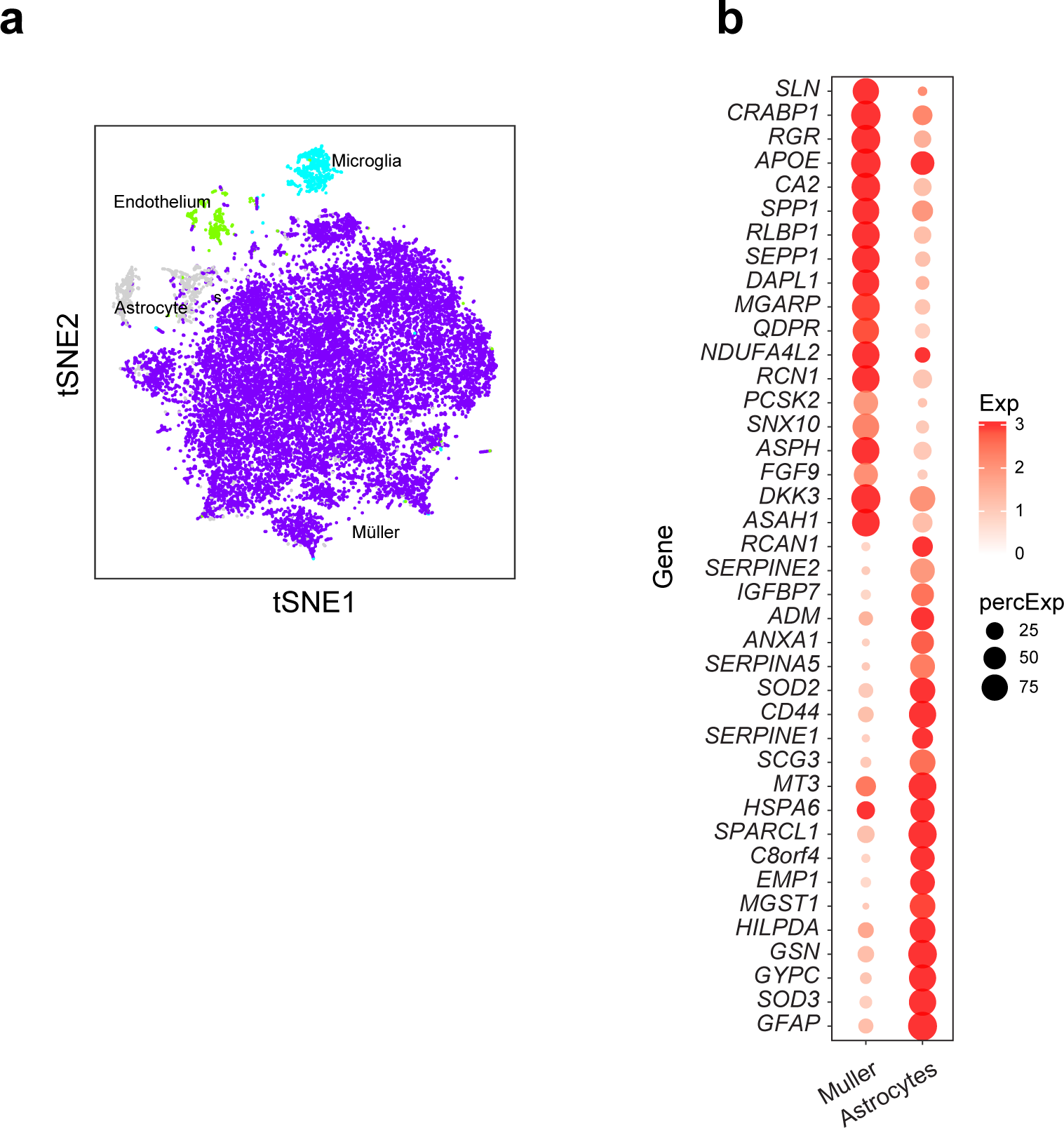
Non-neuronal cell types. (a) Visualization of four non-neuronal types using t-SNE. (b) Genes differentially expressed between Müller glia and astrocytes. Genes shown exhibited >1.5 log fold change.

## Comparison of human and macaque retinal cell types

As noted above, the evolutionary proximity of human and macaque enabled us to name most human clusters based on their striking transcriptional correspondence with types characterized in macaque. We next assessed the extent to which gene expression are conserved among corresponding types between the two species. We compared the expression of type-specific marker genes in 34 corresponding types for which our dataset contained ≥20 cells from each region: 3 photoreceptor, 2 horizontal cell, 12 bipolar cell, 7 amacrine cell, 7 RGC, and 3 non-neuronal types (Fig. 5a-f, see Methods for details). As expected, all corresponding types expressed at least some common type-specific “marker” genes.

**Figure 5.**
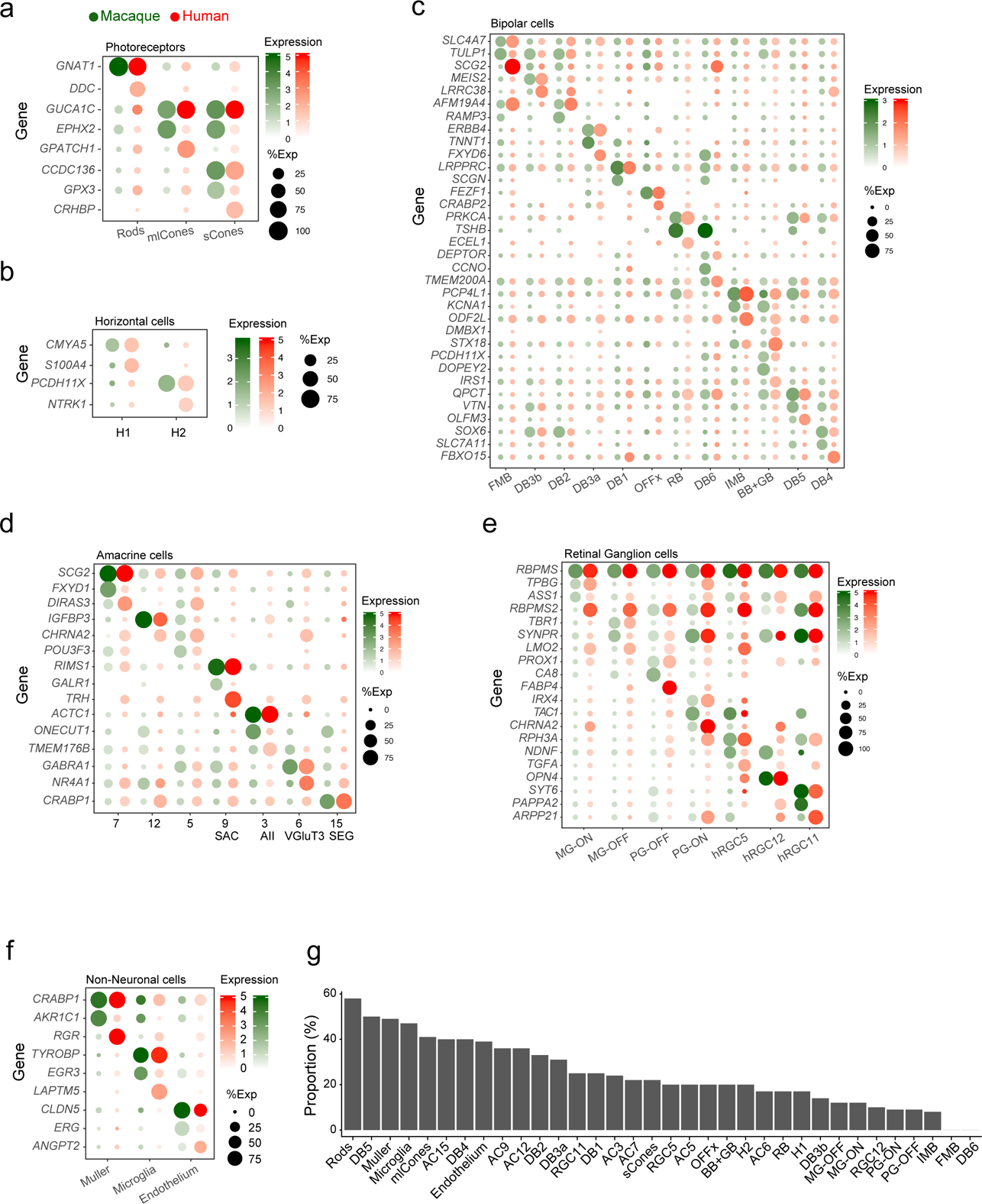
Comparison of gene expression between corresponding human and macaque cell types. (a-f) Dot plots showing similarities and differences in type-specific DE gene expression among corresponding human and macaque types (columns) in PRs (a), HCs (b), BCs (c), ACs (d), RGCs (e), and non-neuronal types (f). (g) Conservation of type-specific marker genes between human and macaque. Graph shows the proportion of DE genes in human (log fold change > 0.5 and adjusted p<0.001 for each type compared to all other types within the class) that are also DE in macaque within shared types (columns).

In some cases, however, type-specific genes were expressed selectively and at high levels in only one of the two species. Examples of DE genes include: (a) *EPHX2* by macaque but not human cones; (b) *GPATCH1* and *CRHBP* by human but not macaque cones; (c) *CA8* by macaque but not human OFF parasol RGCs; (d) *FABP4* by human but not macaque OFF parasol RGCs; (e) *SCGN* by macaque but not human bipolar types DB1 and DB6; (f) *RBPMS2* by human but not macaque midget RGCs; and (g) *RGR* by human but not macaque Müller glia (Fig. 5a-f). As another metric of similarity, we identified genes differentially expressed by each shared human and macaque type (log fold change ≥0.5, <0.001 adjusted p value for each type compared to other types within the class). We then calculated the proportion of DE genes in human that were also DE genes in macaque. The five pairs with the largest proportion of shared DE genes were 2 photoreceptor types, 2 non-neuronal types, and one bipolar type (Fig. 5g).

We used histological methods to validate some of these differences. Labeling with anti-secretagogin (SCGN) plus anti-VSX2 (CHX10), which labels all bipolar cells, confirmed that SCGN is expressed by both bipolar and amacrine cells in macaque retina but only by amacrine cells in human retina (Fig. 6a). Conversely, *RBPMS2* was expressed by human but not macaque midget RGCs, while the canonical markers, *RBPMS* and *SLC17A6* (VGLUT2) were expressed by most or all RGCs in both species (Fig. 6b).

**Figure 6.**
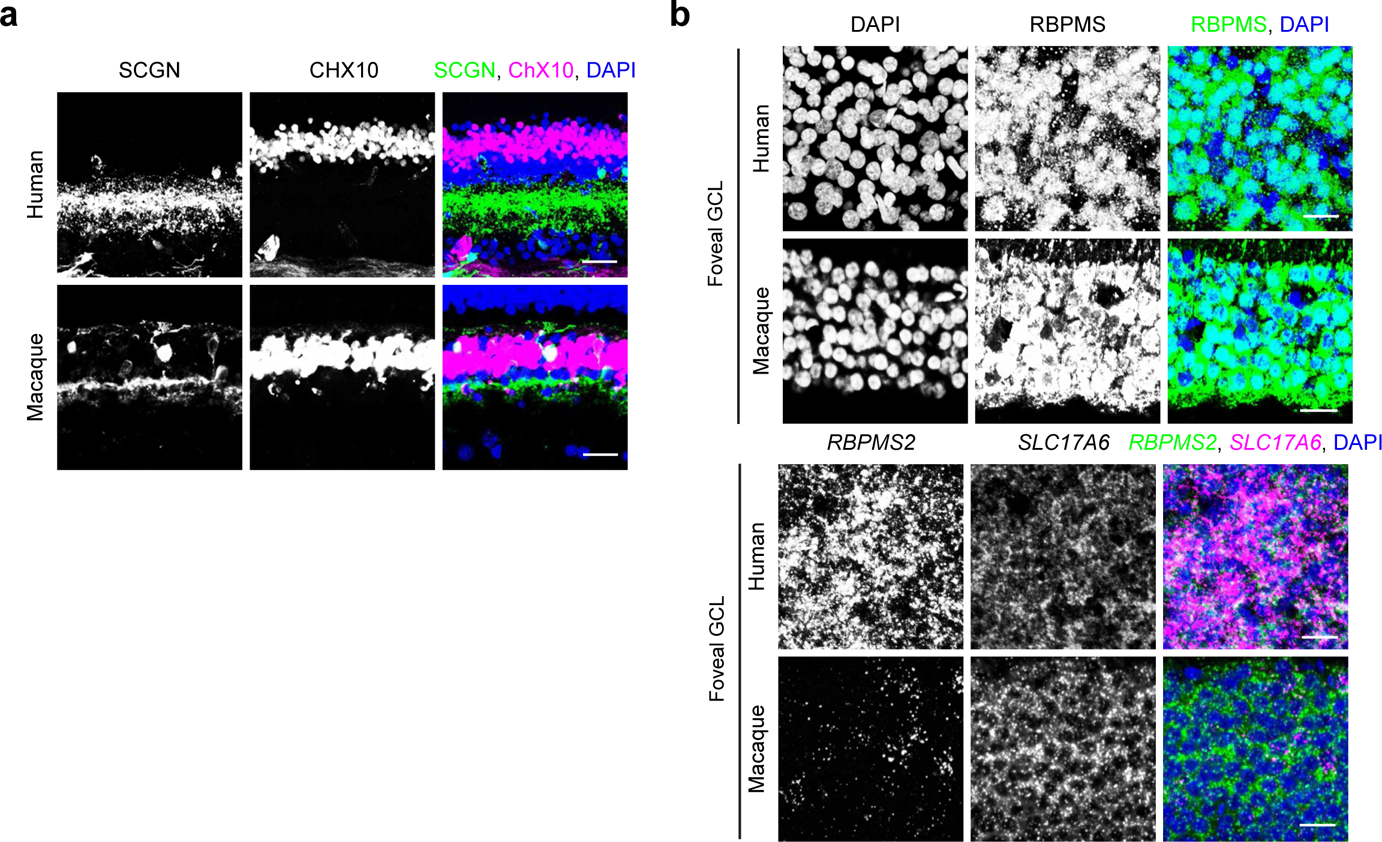
Histological validation of gene expression difference between human and macaque. (a) In human retina (top), SCGN (green) does not label any CHX10-positive BCs (magenta), while SCGN (green) labels several BC (CHX10-positive, magenta) types in macaque retina (bottom). Anti-SCGN also labels amacrine cells in both species. (b) RBPMS detected immunohistochemically (top two rows) is similar in human and macaque GCL, while the expression of RBPMS2 detected by *in situ* hybridization (bottom two rows) is unique to human RGCs, but not found in macaque RGCs. Human RGCs are labeled with *SLC17A6* (magenta). Scale bars are 20 μm.

Together, these comparisons demonstrate predominant but not complete conservation of gene expression by corresponding cell types in human and macaque retina.

## Comparison of fovea and peripheral retinal cells

For analyses presented so far, we pooled data from fovea and periphery. We next compared the regions with each other. Nearly all (57/58) cell types were present in both regions. One GABAergic amacrine type, C18, was found only in the fovea.

For all corresponding types, however, some genes were differentially expressed between foveal and peripheral cohorts. Of 47 types for which there were enough cells in both regions (>20) to enable a comparison, the number of differentially expressed genes ranged from 5 to 100 (log fold change >1; adjusted p-value <0.001). The types with the most differences by these criteria were RGCs (5 types), non-neuronal cells (3 types) and M/L cones (Fig. 7a). Examples include *EPB41L2* and *VTN* expressed by foveal, but not peripheral cones; *TTR* expressed at higher levels by foveal than peripheral bipolar type DB3b and DB4; *TULP1* expressed by peripheral but not foveal bipolar type FMB and DB2; and *RND3* expressed by peripheral but not foveal ON parasol RGCs (Fig. 7b).

**Figure 7.**
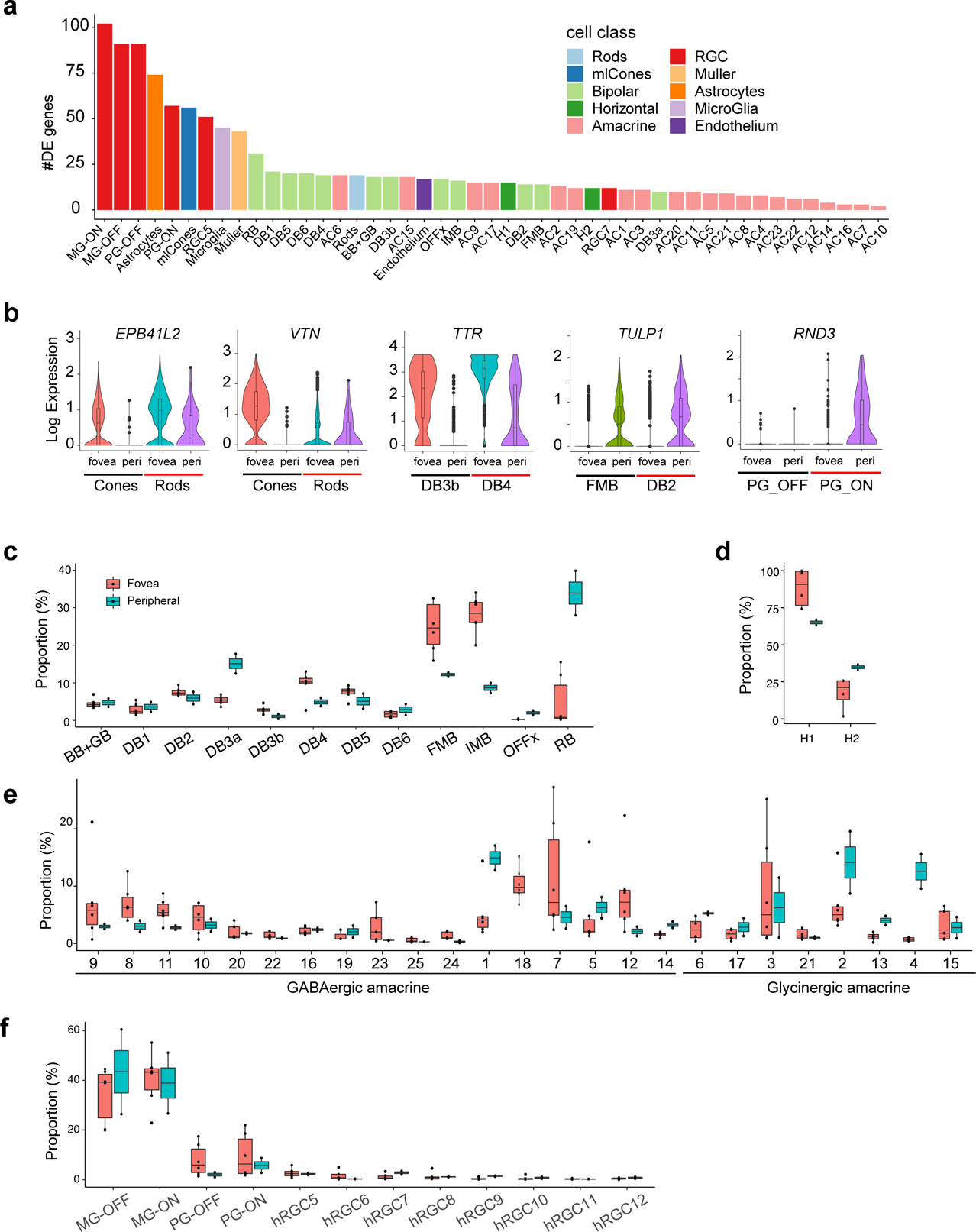
Differences between corresponding cell types in fovea and periphery. (a) Bar plot showing the number of differential expression (DE) genes (log fold change>1 and adjusted p value < 0.001) per matched cell types between fovea and periphery (x-axis). (b) Violin plots showing the expression of select DE genes in relevant foveal and peripheral cell types. (c-f) Box-and-whisker plots showing proportions of cell types in fovea and peripheral retina for BCs (c), HCs (d), ACs(e), RGCs (f). Black horizontal line, median; bars, interquartile range; vertical lines, minimum and maximum, dots are values from individual retina samples, sample size n=6 in fovea, and n=2 in periphery.

In many cases, proportions of cell types also differed between fovea and periphery. Several differences were consistent with previous reports, such as the relatively lower proportion of S cones among all cones in the fovea compared to the periphery (28, 29); the depletion of astrocytes from fovea (30, 31) (0.9% of all non-neuronal cells in fovea and 12% in periphery); and the higher ratio of cone bipolar to rod bipolar cells in the fovea compared to the periphery (32) (rod BCs 35% of peripheral BCs vs 3% of foveal BCs) Fig. 7c). Other differences have not, to our knowledge, been noted previously.

The H1:H2 ratio was nearly ∼4-fold higher in the fovea (7.3:1) than in peripheral retina, 1.9:1; Fig. 7d). The ratio of GABAergic to Glycinergic AC types was higher in fovea (1.8:1) than in the periphery (1.1:1). Several AC clusters showed enrichment in either fovea (e.g., C8, 12 and 23) or peripheral retina (e.g., C1, 2, and 4) (Fig. 7e). The OFF parasol RGC is the only type enriched in fovea using the same criteria, while 5 out of the 8 less abundant RGC types (hRGC cluster 7, 8, 9, 10, 12) were more abundant in peripheral retina than in fovea (Fig. 7f). Foveal enrichment of H1 horizontal cells and OFF parasol RGCs was also observed in the cynomolgus macaque (12).

## Expression of genes implicated in retinal disease

We used the cell atlas to assess retinal expression of 1,756 genes associated with diseases in which vision loss results primarily from death or dysfunction of retinal cells. They include retinitis pigmentosa, cone-rod dystrophy, Leber congenital amaurosis, congenital stationary night blindness, hereditary maculopathy, Leber hereditary optic neuropathy, dominant optic atrophy, open angle glaucoma, age-related macular degeneration, diabetic retinopathy, diabetic macular edema and Macular Telangiectasia type 2. Of these, 624 genes showed robust expression (detected in more than 20% cells of any class, foveal or peripheral, with average expression level > 0.5). We evaluated these, as well as some that fell below threshold but are clinically interesting, further.

We assessed expression of these genes in 9 cell classes: rods, cones, HCs, BCs, ACs, RGCs, Müller glia, astrocytes, microglia and endothelial cells. All genes are listed in Fig. S2, and examples are shown in Fig. 8. Note that the retinal pigment epithelium, which plays a critical role in the pathogenesis of many retinal diseases was not included in our atlas (see Discussion). We summarize these groups here, beginning with diseases for which monogenic high penetrance causes have been identified. We then discuss disorders for which few monogenic causes are known, but numerous susceptibility factors have been implicated primarily through genome-wide association studies (GWAS).

**Figure 8.**
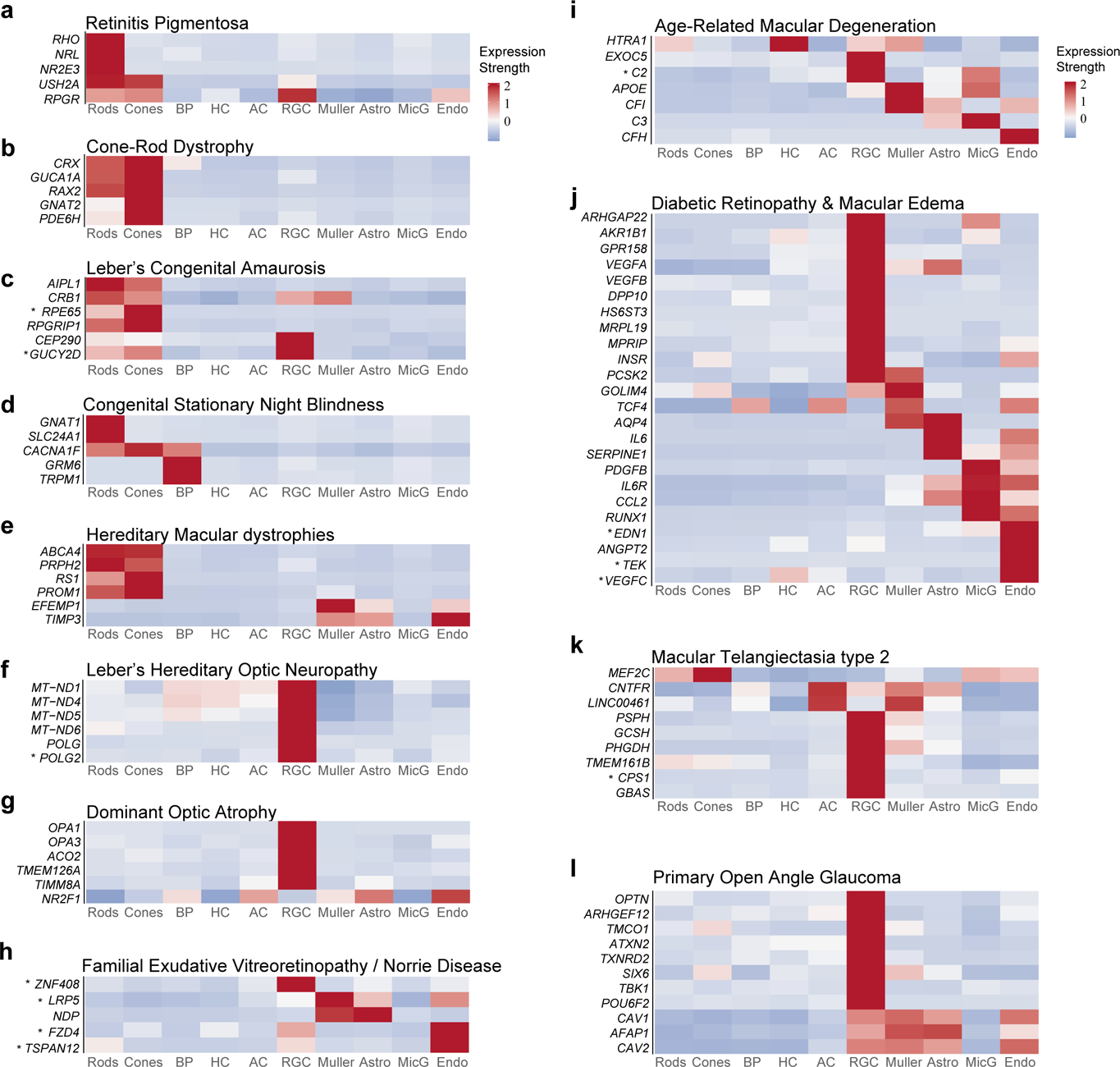
Expression of disease-related genes in retinal cell classes. (a-l) Heat maps show expression of genes implicated in each of 12 groups of blinding diseases described in the text. Color represents the scaled expression level of genes among all cell classes. Thus, the heat maps accurately reflect the order of expression among cell classes for each gene but not the absolute levels of expression. Asterisks mark genes that were included because of clinical interest but fell below the threshold noted to the text.

Retinitis Pigmentosa (RP) is a diffuse photoreceptor dystrophy that also affects the pigment epithelium. It manifests as night blindness with progressive visual field loss(33). Clinical features in the macula often include loss of foveal reflex, abnormalities at the vitreoretinal interface, and cystoid macular edema. Other typical findings include arteriolar narrowing, waxy pallor to the optic disc, and variable amounts of bone-spicule pigment changes. Consistent with the predominant functional deficits of night blindness, most genes implicated as monogenic causes of RP were predominantly expressed in rods (Fig. 8a, S3). Potentially consistent with other clinical findings of RP, some genes were also expressed in vascular endothelium and RGCs (e.g. *RPGR* and *TOPORS)*, while others were expressed at highest levels in RGCs (e.g. *SLC25A46*, *SLC7A14* and *RP9*) or Müller glia (e.g. *RGR* and *RLBP1)*. As noted above, some genes in this and other disease groups (e.g., *RPE65*) are likely to act in and be expressed at higher levels by retinal pigment epithelial cells, which were not included in our dataset.

Cone-rod dystrophy affects both photoreceptor classes and patients with this condition demonstrate expanding central scotomas often leading to severe visual impairment (34). Consistent with this pathology, causative genes were expressed in both rods and cones (e.g., *CRX*, *RAX2*), with expression often higher in the latter (e.g., *GNAT2, PDE6H*) (Fig. 8b).

Lebers Congenital Amaurosis (LCA) is a severe group of inherited retinal dystrophies characterized by nystagmus, sluggish or absent pupillary light reflexes and blindness, often in the first year of life (35). Genes mutated in the most prevalent forms of LCA were expressed in both rod and cone photoreceptors, consistent with the characteristic early absence of retina-wide rod and cone photoreceptor function demonstrable by electroretinogram (ERG). Several (*CEP290, GUCY2D and CRB1*) were also expressed in RGCs (Fig. 8c).

Congenital Stationary Night Blindness (CSNB), a lifelong, nonprogressive abnormality of scotopic vision, disrupts transmission through the rod pathway by disabling neurotransmission from rods to rod bipolar cells. In most cases, cell loss is minimal.

Genes implicated in CSNB were generally expressed either in rods (e.g. *GNAT1, SLC24A1*) or bipolar cells (e.g. *GRM6, TRPM1*). *CACNA1F*, which harbors mutations in the majority of cases of incomplete X-linked CSNB, was expressed in both photoreceptor and bipolar cells, consistent with its wider phenotypic spectrum encompassing X-linked progressive cone-rod dystrophy, optic atrophy, and Åland Island eye disease (AIED) (Fig. 8d). Finally, although, *NYX* and *LRIT3*, which harbor CSNB mutations (36, 37) were expressed at levels too low to meet our set screening threshold, they were preferentially expressed in bipolar cells and photoreceptors, respectively.

Macular dystrophies, including Stargardt Disease, Vitelliform degenerations, Pattern Dystrophies, Sorsby Macular Dystrophy and Familial drusen, are slowly progressive retinal degenerations that account for a significant proportion of cases of central vision loss among adults under the age of 50 (38). Genes involved in these dystrophies were generally enriched in photoreceptors (e.g. *PRPH2*, *PROM1*), but others were also expressed in non-neural cells – e.g. *EFEMP1* selectively in Müller Glia and *TIMP3* selectively in vascular endothelium (Fig. 8e). *TIMP3,* which is mutated in Sorsby Macular Dystrophy, encodes a protein involved in matrix remodeling and suppression retinal angiogenesis. Although the primary site of pathogenesis in this disease is believed to be at the level of retinal pigment epithelium or Bruch’s membrane, our results suggest that might also act within retinal vasculature.

Leber’s hereditary optic neuropathy (LHON) is the most common inherited mitochondrial disease with ophthalmic manifestations. It is caused by mutations in genes primarily encoding respiratory complex chain 1 proteins (e.g. *ND1, ND4, ND6*), leading to defects in NADH-ubiquinone oxidoreductase chains that may impair glutamate transport and increase production of reactive oxygen species (39). The result of these impairments is RGC dysfunction and, eventually, apoptosis, and atrophy of the retinal nerve fiber layer. Consistent with this pathogenesis, all LHON genes were predominantly expressed in RGCs (Fig. 8f).

Autosomal Dominant Optic Atrophy, the most common hereditary optic neuropathy, is characterized by gradual loss of visual acuity that is generally bilateral and symmetric. RGC degeneration, particularly in the papillomacular bundle, has been implicated as the primary mechanism of disease (40). Consistent with this pattern of expression, causative genes such as *OPA1* and *OPA3* were predominantly expressed in RGCs (Fig. 8g).

Inherited Vitreoretinopathies include Familial Exudative Vitreoretinopathy (FEVR), Norrie Disease and Coats Disease. Most genes mutated in these disorders were expressed in retinal vascular endothelium (Fig. 8h). In addition, however, *NDP* and *LRP5* were also expressed in Müller glia (41) and in astrocytes, a finding compatible with observations of *NDP* expression in a subset of cortical astrocytes (42)

Age Related Macular Degeneration (AMD) is broadly classified into non-exudative (“dry”) and exudative or neovascular (“wet”) types. In dry AMD mild forms are characterized by drusen accumulation between retinal pigment epithelium and Bruch’s membrane, which can progress to late forms with large patches of atrophic outer retina. In wet AMD, aberrant angiogenesis originating either within the choroid or retina leads to often catastrophic sub- or intraretinal hemorrhage (43). *HTRA1*, a major susceptibility gene for neovascular AMD (5), was expressed at high levels in HCs and Müller glia (Fig. 8i). While the disease-related effects of this gene are thought be exerted in the retinal pigment epithelium, its expression in HCs and Müller glia suggests additional sites of action. Alleles in the *CFH* gene and other complement pathway genes have also emerged as risk factors for AMD; we noted expression of multiple complement genes (*CFH, CFI, C2, C3*) in different cell classes (Fig. 8i).

Diabetic retinopathy (DR) and Diabetic Macular Edema (DME) together represent the leading cause of blindness and visual disability among working-age adults of all races living in industrialized nations (44). While many genes implicated in diabetic retinopathy – classically considered a predominantly microvascular complication of diabetes mellitus – were expressed in non-neuronal retinal cell classes (particularly, vascular endothelial cells), a large proportion, such as *HS6ST3* (45), DPP10 (46), and *VEGFB,* were almost exclusively expressed in RGCs (Fig. 8j).

Macular Telangiectasia type 2 (Mac Tel 2) is a rare retinal neurodegenerative condition that leads to late-onset progressive central vision loss (47, 48). Early clinical findings, including retinal discoloration and capillary telangiectasis, are limited to the perifoveal region. Photoreceptor loss and foveal atrophy occur as the disease progresses. Mac Tel 2 is currently considered to be a primary neurodegenerative condition of the retina with secondary vascular involvement rather than (as previously hypothesized) a primary vasculopathy. Several risk alleles and genes in proximity to SNPs identified by GWAS studies implicate Müller cell dysfunction and dysregulation of serine metabolism in (49, 50) pathogenesis. We found expression patterns compatible with these hypotheses. For example, *PHGDH* and *PSPH*, encoding enzymes involved in L-serine synthesis, were selectively expressed in RGCs with *PHGDH* also expressed in Müller glia. Finally, to gain insight into the protective effects of CNTF, which is currently in phase 2 clinical trials for Mac Tel 2, we investigated the expression of CNTFR. We found expression in RGCs and Müller glia as well as amacrine cells and astrocytes (Fig. 8k).

Open Angle Glaucoma, a primary optic neuropathy characterized by loss of RGCs, is one of the most common causes of vision loss world-wide and the leading cause of irreversible blindness among African Americans (51, 52). Among genes implicated in glaucoma, either by GWAS or as rare Mendelian alleles, most were expressed predominantly within RGCs (e.g., *OPTN, TMCO1, TBK1)* (Fig. 8l). Other genes, including *FOXC1, CYP1B1, LMX1B* and *MYOC* were not expressed substantially in any neural retina cell classes, consistent with their proposed role predominantly in the anterior segment with mutations leading to high intraocular pressure, a major risk factor for glaucoma (Fig. S3). Indeed, in a parallel study, we have demonstrated expression of these genes in cells of the aqueous humor outflow pathways (53).

In general, these patterns of expression match those we previously documented for macaque. For example, of the 85 genes shown in Fig. 8, 79 were profiled in macaque and of these, 73 (or 92%) were expressed at highest levels in the same cell class in both species.

We next compared foveal and peripheral cohorts of cell classes in which genes were highly expressed (Fig. 9a and 9b). Several patterns were consistent with clinical features of the associated conditions. For example, many genes with causative mutations leading to Retinitis Pigmentosa, including *RHO*, *NRL*, and *NR2E3* demonstrated foveal rod enrichment (Fig. 9a and S3). *RP1* was preferentially expressed in foveal rods and cones. In contrast, *PDE6H,* a cone-rod dystrophy gene, demonstrated preferential expression in peripheral rods. Genes implicated in macular dystrophies were generally enriched in the fovea compared to the peripheral retina.

**Figure 9.**
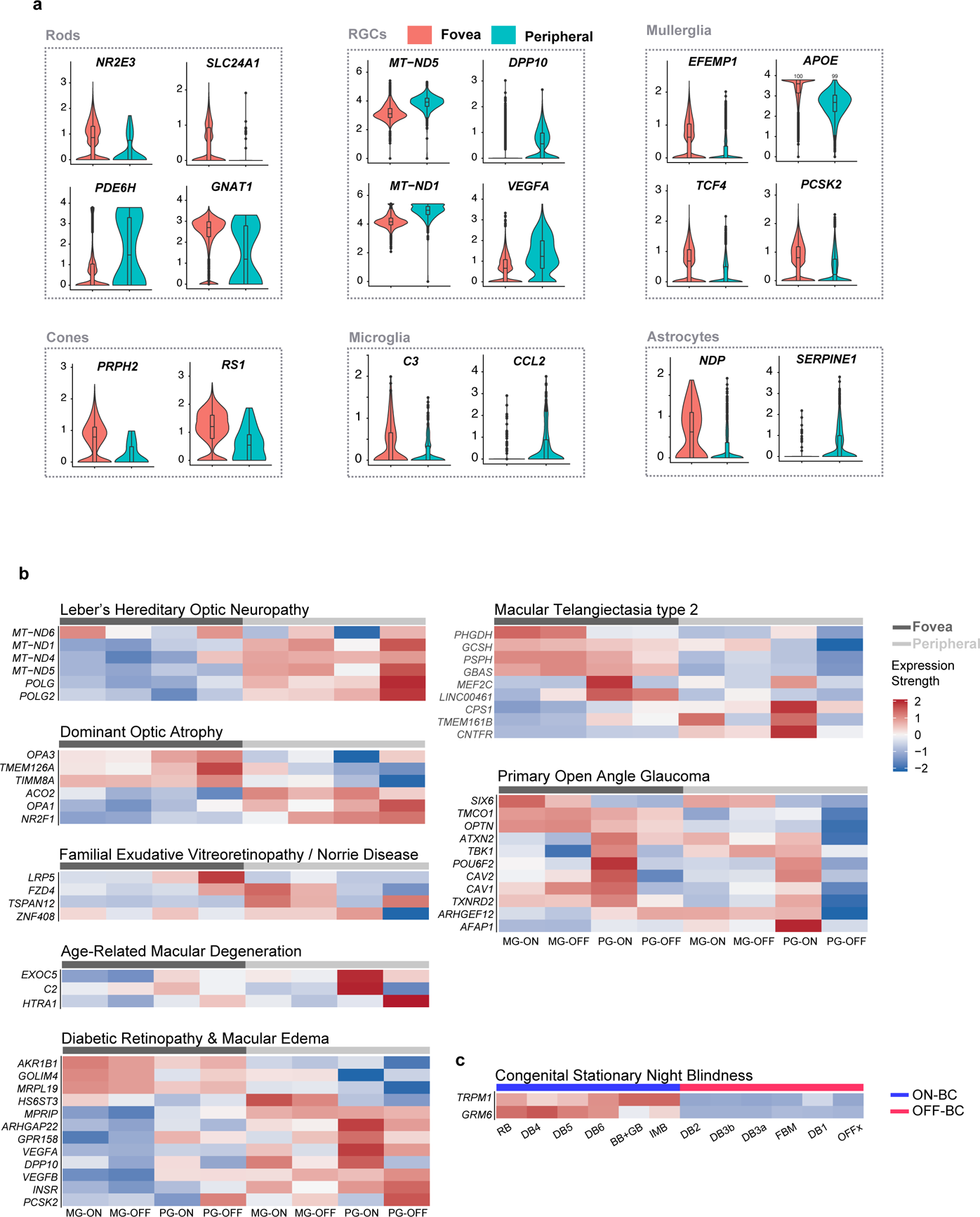
Differential expression of disease-related genes by region and cell type. (a) Violin and superimposed box plots showing differential expression of select disease genes between foveal and peripheral cell classes. (b) Heat maps showing expressions of select disease genes in four major RGC types. Cell types are segregated by their regions: fovea versus periphery. (c) Heat map showing expression of *GRM6* and *TRPM1*, two previously reported congenital stationary night blindness genes, in ON bipolar cells (blue bar, top), but not in OFF bipolar cells (red bar, top)

*ABCA4*, which harbors causative mutations leading to Stargardt disease, was enriched in foveal photoreceptors (Fig. S3); *EFEMP1*, implicated in Doyne Honeycomb Dystrophy, was predominantly expressed in foveal Müller glia. *APOE*, implicated in Open Angle Glaucoma, as well as *PHGDH* and *CNTFR* were expressed at higher levels in foveal than peripheral Müller glia. In contrast, *VEGFA*, polymorphisms of which have been linked to severity of Diabetic Retinopathy, was expressed at higher levels in the periphery compared to fovea.

Finally, we assessed type-specific expression in RGCs for genes implicated in dominant optic atrophy, diabetic retinopathy, diabetic macular edema, Mac Tel 2 and primary open angle glaucoma, and type-specific expression in bipolar cells for genes implicated in CSNB (Fig. 9b, c). Genes with type-specific RGC expression patterns included the glaucoma-associated genes *SIX6*, which was enriched in midget ganglion cells; OPTN, enriched in foveal RGCs; *CAV2* and *POU6F2* enriched in ON parasol RGCs and *AFAP1*, enriched in peripheral ON parasol RGCs. (*POU6F2* was expressed at highest levels in RGC types 5, 11 and 12, which include ipRGCs; data not shown.) Several but not all genes implicated in Mac Tel 2 (e.g. *PHGDH*, *PSPH*, *LINC00461* and *GBAS*) were enriched in foveal RGCs. Patterns of expression differed, however, with *PHGDH* expressed primarily in foveal midget RGCS, *LINC00461* primarily in foveal parasol RGCs, and *PSPG* and *GBAS* in both. *MRPL19*, one of the few genes implicated specifically in DME, which affects the fovea by clinical definition, was expressed preferentially in foveal RGCs (54).

## DISCUSSION

We used high-throughput single-cell RNA-seq to generate a cell atlas of the adult human retina. From 55,736 foveal and 29,246 peripheral retinal cells, we identified 58 cell types. We then used the cell atlas to compare fovea with peripheral retina, and human with macaque retinal cell types. Finally, we probed region-, cell class-, and cell type-specific expression of genes associated with blinding retinal diseases. Together, our atlas provides a roadmap for human retinal research and paves the way for further research on the pathology of ocular diseases.

## Human cell atlas

Non-diseased retina is seldom excised during ocular surgery, so tissue must be obtained postmortem. Given that cell viability declines and transcriptomic profiles change after death, with dramatic alterations after 10 hours post-mortem, data quality hinges in large part on the time between death and tissue processing. For example, Lukowski et al. (55) showed that rods began to degenerate and their expression of *MALAT1*—a long non-coding RNA—decreased at this point. In our dataset, retinas from 6 of the 7 donors were obtained within 6.5hr post-mortem, all rod photoreceptors clustered together, and *MALAT*1 levels were high. These results affirm the high quality of the cells from which the atlas was generated.

We and others have recently reported results of scRNA-seq studies on human retina (55–60) (see Table S2). However, our initial study was focused on bipolar cells, and some groups used fetal rather than adult cells, and/or did not distinguish foveal from peripheral cells. Three of these groups, however, used adult retina and separated fovea from peripheral retina. Although these studies generated valuable data, they were disadvantaged in that rods comprise a large fraction of all cells (>70%), reducing power to distinguish cell types among less abundant classes. This problem is most severe for RGCs, which comprise <2% of retinal cells. We circumvented these limitations by depleting rods in some samples (using anti-CD73) to enrich other neuronal classes, and by selecting RGCs in other samples (using anti-CD90). These strategies allowed us to distinguish more types within classes than in previous studies. For example, we were able to characterize vGlut3 excitatory ACs (0.7% of total retinal cells), ipRGCs (0.02% of total retinal cells), S cones (0.07% of total retinal cells), and the primate-specific OFFx bipolar type (12). Thus, our cell atlas represents the most complete classification to date of cell types in adult human retina.

## Comparison with monkey atlas

A major hurdle for studying human retinal biology and diseases is that retinas of accessible animal models differ from human retina in critical respects. Perhaps most important is that among mammals, only primates have a fovea or macula. The fovea comprises only ∼1% of retinal area in humans, but accounts for most of our high-acuity vision, much of our chromatic vision, and supplies ∼50% of the visual input to the cortex (61, 62). Moreover, the fovea, and the macula within which it is embedded, are the principal site of pathology among diseases such as age-related macular degeneration, diabetic macular edema, hereditary macular dystrophy, and macular telangiectasia.

Lacking fovea and macula, it is unsurprising that rodent models of these diseases have severe limitations.

Our results address this issue in two ways. First, by generating a human cell atlas, and a comprehensive database on expression of disease-related genes, we provide a foundation for both translational and basic studies. Second, by documenting close similarities between human retinal cell types and the macaque types we described recently (12), we both validate the use of this non-human primate model and point out some important differences that will need to be considered in interpreting studies of non-human primates. Although some differences could result from imperfections in gene and transcript annotation, it is likely that the vast majority are genuine.

## Difference between the fovea and peripheral retina

The structural and functional difference between the fovea and peripheral retina could result from the existence of specialized foveal cell types. We show however, that nearly all retinal cell types are shared between fovea and periphery in human retina. There are, however, substantial regional differences in gene expression and abundance between foveal and peripheral cohorts of shared types. Limitations to the comparison include low cell number in periphery for some cell types, and potential bias introduced by the methods we used to deplete rods and enrich RGCs from peripheral samples.

Nonetheless, many of the differences in abundance we observed were consistent with those reported by others based on morphological analysis, and we reported histological validation of some of the DE genes in a recent study (12). Thus, in humans as in macaques, the fovea and peripheral retina are composed of similar cell types, with the structural and functional differences between them likely arising from differences in abundance of shared types and specific aspects of gene expression.

## Mapping disease genes to cell classes and types

We analyzed expression of 636 genes associated with retinal diseases, chosen from an initial list of 1,756. Although long, the list is incomplete, because retinal pigment epithelium, which plays a major role in the pathogenesis of retinitis pigmentosa and age-related macular degeneration, was not included in our dataset. Moreover, our criterion for inclusion was expression in more than 20% of cells of at least one class in either fovea or peripheral retina, so genes expressed at slightly lower levels or in only a few minor types within a class would have been excluded. We added 12 genes that fell below threshold to the 624 that met the criterion based on their known clinical relevance, but others remain to be analyzed.

We document cell-class and cell-type specific expression patterns for many of these genes. While expression patterns for many of them supported prior reports of pathogenetic mechanisms (detailed in Results), others provided unexpected insights into potential cellular contributors to disease. For example, mutations responsible for many genes implicated in retinitis pigmentosa – in which waxy pallor of the optic disc and visual field loss are common clinical findings – demonstrated selective expression not only by PRs but also by RGCs. Similarly, many genes implicated in diabetic retinopathy – classically considered a retinal vasculopathy– were expressed at relatively high levels by RGCs, suggesting that a primary neuropathic process may also be involved. In some cases, we identified RGC type specificity among these genes. Of particular interest are genes implicated in POAG, given the current lack of knowledge about whether specific RGC types exhibit selective vulnerability in this disease (63).

Together, our results offer new insights into many rare and common retinal diseases, and may contribute to a more comprehensive understanding of their pathogenesis and the uncovering of novel therapeutic targets.

## MATERIALS AND METHODS

### Human tissue

Human eyes used for sequencing and histological studies were collected 3-14 hours post mortem through the Rapid Autopsy Program, Massachusetts General Hospital, with all but one collected ≤6.5 hours post mortem (Table S1). The globe was immediately transported back to the lab in a humid chamber. Hemisection was performed to remove the anterior chamber, and the posterior pole was immersed in Ames equilibrated with 95% O2/5% CO2 before further dissection and dissociation. All donors were confirmed to have no history or clinical evidence of ocular disease or intraocular surgery.

For histological studies, dissected retinal tissues were fixed in ice-cold 4% PFA for 2hrs. Acquisition and use of human tissue was approved by the Human Study Subject Committees (DFCI Protocol Number: 13-416 and MEE - NHSR Protocol Number 18-034H). For transcriptomic analysis, retinal cells were dissociated as described in the next section.

## Single cell isolation, library preparation and sequencing

Single cell libraries were generated by minor modifications of methods developed for macaque retina (12). Briefly, a ∼1.5 mm diameter circular region centered on the foveal pit was dissected from the retina, and peripheral retinal pieces were pooled from all retinal quadrants. Dissected tissues were digested with papain (Worthington, LS003126) for 30min at 37°C. Following digestion, samples were dissociated and triturated into single cell suspensions with 0.04% bovine serum albumin (BSA) in Ames. Dissociated cells from digested peripheral retinas were incubated with CD90 microbeads (Miltenyi Biotec, 130-096-253; 1 ml per 10^7^ cells) to enrich RGCs or with anti-CD73 (BD Biosciences, clone AD2; 5 ml per 10^7^ cells) followed by anti-mouse IgG1 microbeads (Miltenyi Biotec, 130-047-102; 10 ml per 10^7^ cells) to deplete rods.

Incubations were at room temperature for 10 min. CD90 positive cells or CD73 negative cells were selected via large cell columns through a MiniMACS Separator (Miltenyi Biotec). Foveal samples were used without further processing. Single cell suspensions were diluted to 500-1800 cells/µL in 0.04% BSA/Ames for loading into 10X Chromium Single Cell v2 or v3 Chips. Following collection, cDNA libraries were prepared following the manufacturer’s protocol, and sequenced on the Illumina HiSeq 2500 (Paired end reads: Read 1, 26bp, Read 2, 98bp).

## Bioinformatics analysis

### Clustering

Sequencing reads were demultiplexed and aligned to a human transcriptomic reference (GRCh38) with the Cell Ranger software (version 2.1.0, 10X Genomics for the v2 samples, and version 3.0.2 for the v3 samples) for each 10X channel separately. The resulting digital gene expression (DGE) matrices representing the transcript counts for each gene (rows) in each cell (columns) were combined for all samples (foveal and peripheral), and analyzed further using the R statistical language following methods described by Peng et al. (12), with minor modifications, as follows.

A threshold of 600 detected genes per cell was applied to filter out low quality cells or debris from the combined DGE matrix. Clustering was performed to first group cells into major cell classes. In addition to the nine cell classes described in the main text, we detected small numbers of epithelial cells and melanocytes (<100 cells); they were not considered further, because of the likelihood that they arise from non-retinal contaminants. For each of the neuronal classes, a second round of clustering was performed to assign cells into types.

The DGE matrix corresponding to each class was separately analyzed to identify molecularly distinct types within that class. First, normalized, log-transformed expression values E_i,j_ for gene i in cell j were calculated following (64). Highly variable genes (HVGs) were identified as in (65), and used for dimensionality reduction. Batch correction was performed using the linear regression approach adapted from the R package Seurat. Principal component analysis (PCA) was performed and statistically significant PCs were estimated using Random Matrix Theory (66). Cell clustering was performed using Louvain algorithm with Jaccard correction (64). Some low quality cells or doublets became apparent only after clustering, so averaged number of genes/transcripts per cell and the proportion of mitochondria gene transcripts were assessed for each cluster.

To visualized the data in 2D space, we applied t-distributed stochastic neighbor embedding (t-SNE), using as input the normalized cell factors computed using the R package ‘liger’(67), which uses an integrative non-negative matrix factorization framework. Averaged expression matrix of HVGs for each cluster were calculated to build dendrogram (Fig. 3b upper panel, hierarchical agglomerative clustering of Euclidean distance metric with complete linkage), which revealed transcriptomic relationships among types. Neighboring clusters on this dendrogram were iteratively merged if no more than five DE gene was found showing ⩾1.1 log fold change with adjusted p value <0.001 using the R package ‘MAST’ (68).

### Comparing human and macaque cell types

We sought correspondence between human and macaque cell types using the multi-class classification approach described earlier (12). Briefly, for each cell class in macaque, we trained a multi-class classifier using the R package ‘xgboost’ (14) to map cells to discrete type labels based on their transcriptional signatures. This classifier was then applied to each human cell of the same class to assign it a macaque type label based on its expression of 1:1 gene orthologs, but in a manner completely agnostic to its cluster identity. Confusion matrices (e.g. Fig. 2b,e,h) were used to identify correspondences between macaque and human types within each class. Shared types were defined as those exhibiting a near 1:1 correspondence using this classification approach.

Each pair of shared human and macaque types were assessed for DE genes compared to other types of the same class (Fig. 5g). Only genes expressed in 20% of cells in either type and exhibiting a ⩾0.5*log fold difference were considered for analysis. The statistic criteria is less stringent comparing to that of the fovea vs peripheral comparison within human cells (next section) to compensate for the across species differences. DE genes were selected as those satisfying a p <0.001 cutoff according to the ‘MAST’ test (56). For each shared type *i* this resuled in two DE lists, one each for human (*hDE*) and *i* macaque (*mDE*) respectively. The proportion of type-specific DE genes that were shared across species (*pDEi*) was computed for each type *i* as,

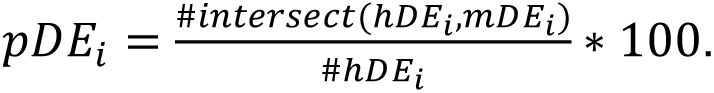

### Comparing foveal and peripheral cells

To evaluate the extent of similarity between foveal and peripheral cell types in human retina (Fig. 7a), we measured the number of DE genes identified between foveal and peripheral cells for each human type. Only types with at least 20 cells in both regions were considered. DE genes in this comparison were those with differences in expression ⩾1*log fold change and adjusted p-value<0.001.

### Immunohistochemistry and fluorescence in situ hybridization

Procedures for tissue preparation, immunohistochemistry and in situ hybridization have been described in (12, 64, 69). Briefly, eyes were fixed in ice-cold 4% paraformaldehyde, rinsed with PBS, immersed in 30% sucrose overnight at 4 °C, embedded in Tissue Freezing Medium (EMS) and cryosectioned at 20 μm. For immunohistochemistry, antibodies were diluted in 3% donkey serum (Jackson, 017-000-121), and 0.3% Triton-X in PBS. Antibodies used for immunostaining were as follows: goat anti anti-CHX10 (1:300, Santa Cruz); rabbit anti-TFAP2A (1:500, DSHB); rabbit anti-Secretagogin (1:10,000; BioVendor). For in situ hybridization, sections were mounted on Superfrost slides (Thermo Scientific), treated with 1.5 mg/mL of proteinase K (NEB, P8107S), and then post-fixed and treated with acetic anhydride for deacetylation. Probe detection was performed with anti-DIG HRP (1:1000) and anti-DNP HRP (1:500), followed by tyramide amplification.

### Image Acquisition and Processing

Images were acquired on Zeiss LSM 710 confocal microscopes with 405, 488-515, 568, and 647 lasers, processed using Zeiss ZEN software suites, and analyzed using ImageJ (NIH). Images were acquired with 16X, 40X or 63X oil lens at the resolution of 1,024×1,024 pixels, a step size of 0.5-1.5µm, and 90µm pinhole size. ImageJ (NIH) software was used to generate maximum intensity projections. Adobe Photoshop CC was used for adjustments to brightness and contrast.

### Mapping disease genes

SNP-trait associations (N=980) were downloaded on 09/03/2019 from the NHGRI-EBI GWAS Catalog (70) for the traits, “open-angle glaucoma” (n=108), “intraocular pressure measurement” (n=504), “diabetic retinopathy” (n=138), “age-related macular degeneration” (n=230) and Macular Telangiectasia Type 2 (n=5). A list of genes and loci associated with retinal diseases was downloaded on 09/03/2019 from the Retinal Information Network, last updated on 07/01/2019 (RetNet, https://sph.uth.edu/retnet/).

## Data availability

The accession number of raw and processed single cell RNAseq data reported in this study is GEO: XXXX (in process). Data can be visualized at the Broad Institute’s Single Cell Portal: https://singlecell.broadinstitute.org/single_cell/.

## ACKNOWLEDGEMENTS

We thank the donors and their families for their generous and essential role in this research. This work was supported by National Institutes of Health grants K12 EY016335-13, R21 EY028633, K99 EY028625 and U01 MH105960 and a Chan Zuckerberg Initiative Human Cell Atlas grant CZF2019-002459.

