## Supplemental figures and tables for "Cell Atlas of the Human Fovea and Peripheral Retina"

**Figure S1**

tSNE visualization showing contributions to cell types by batch for photoreceptors (a), horizontal cells (b), bipolar cells (c), amacrine cells (d), retinal ganglion cells (e) and non-neuronal cells (f). Each dot represents one cell. Colors distinguish retina samples. Source of each sample is shown in Table S1. Overall, batch effects were minimal.

**Figure S2**

Violin and superimposed box plots showing expression of OPN4 in RGC clusters

**Figure S3**

Heat maps showing expression patterns of disease genes by cell classes in the fovea and periphery. Only genes expressed by more than 20% of cells in any individual class in either fovea or peripheral cells are plotted.

**Table S1**

Information on donors from whom retinal cells were obtained for scRNA-seq profiling.

**Table S2**

Publications reporting single cell or single nucleus profiling on cells from human retina.

Figure S1

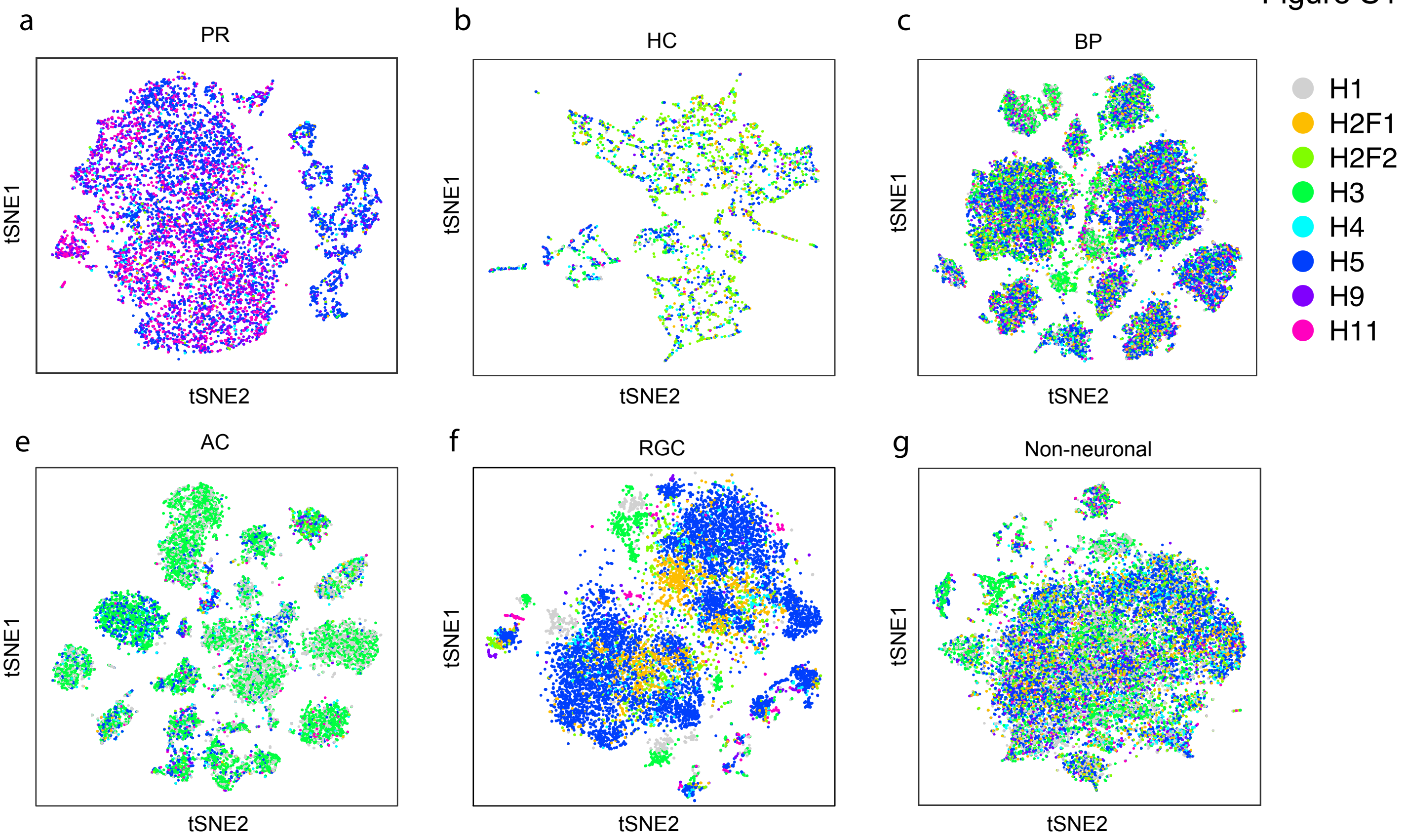

a
PR
tSNE1
tSNE2
b
HC
tSNE1
tSNE2
c
BP
tSNE1
tSNE2
H1
H2F1
H2F2
H3
H4
H5
H9
H11
e
AC
tSNE1
tSNE2
f
RGC
tSNE1
tSNE2
g
Non-neuronal
tSNE1
tSNE2

Figure S2

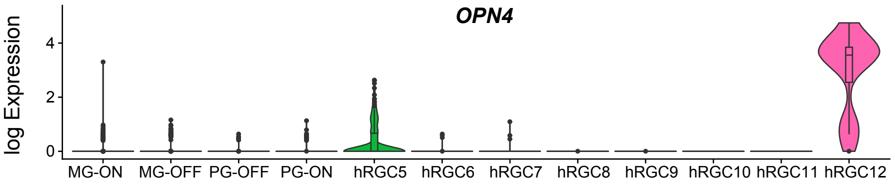

OPN4
log Expression
4
2
0
MG-ON
MG-OFF
PG-OFF
PG-ON
hRGC5
hRGC6
hRGC7
hRGC8
hRGC9
hRGC10
hRGC11
hRGC12

### Figure S3

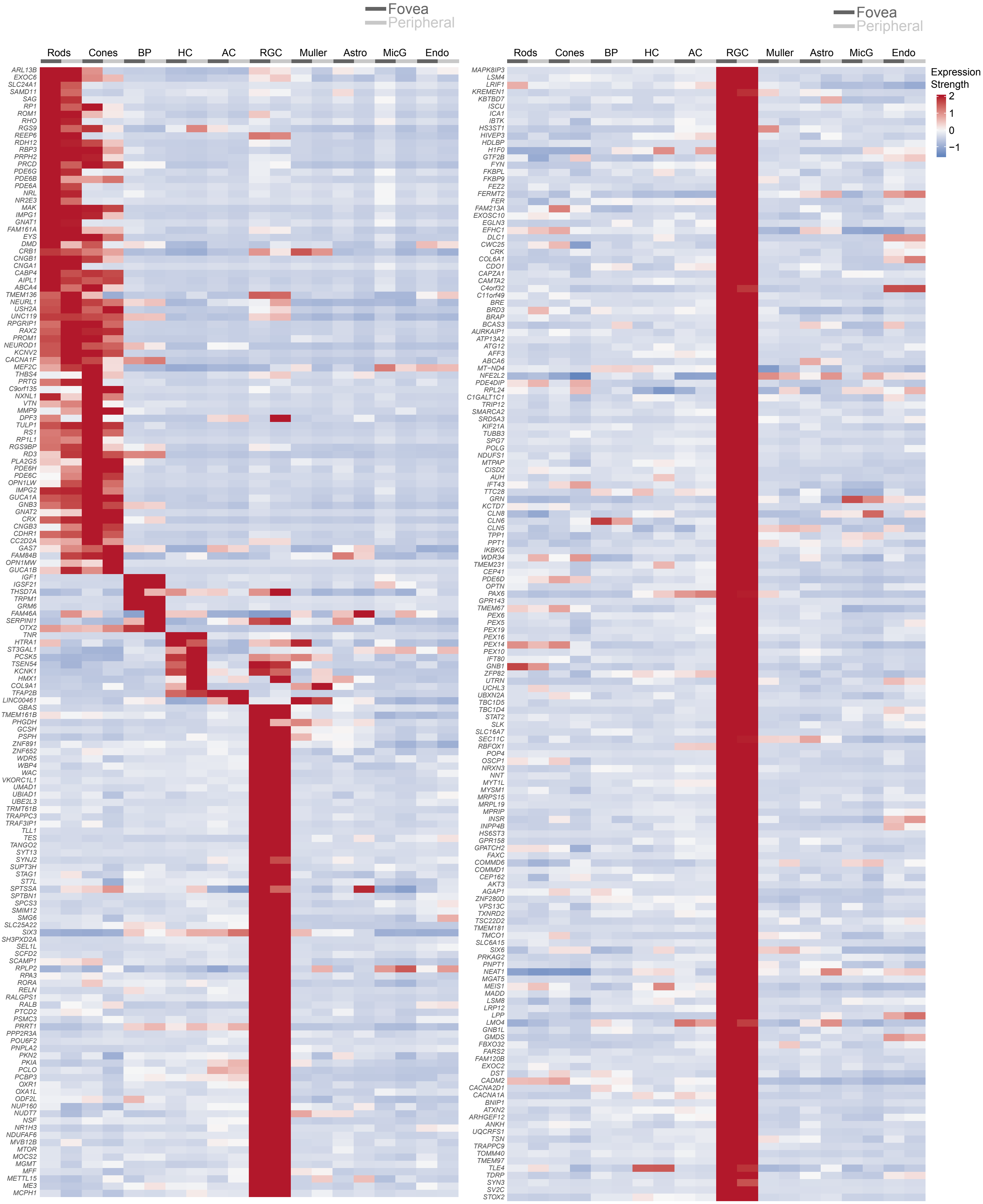
Fovea
Peripheral
Rods
Cones
BP
HC
AC
RGC
Muller
Astro
MicG
Endo
ARL13B
EXOC6
SLC24A1
SAMD11
SAG
RP1
ROM1
RHO
RGS9
REEP6
RDH12
RBP3
PRPH2
PRCD
PDE6G
PDE6B
PDE6A
NRL
NR2E3
MAK
IMPG1
GNAT1
FAM161A
EYS
DMD
CRB1
CNGB1
CNGA1
CABP4
AIPL1
ABCA4
TMEM136
NEURL1
USH2A
UNC119
RPGRIP1
RAX2
PROM1
NEUROD1
KCNV2
CACNA1F
MEF2C
THBS4
PRTG
C9orf135
NXNL1
VTN
MMP9
DPF3
TULP1
RS1
RP1L1
RGS9BP
RD3
PLA2G5
PDE6H
PDE6C
OPN1LW
IMPG2
GUCA1A
GNB3
GNAT2
CRX
CNGB3
CDHR1
CC2D2A
GAS7
FAM84B
OPN1MW
GUCA1B
IGF1
IGSF21
THSD7A
TRPM1
GRM6
FAM46A
SERPINI1
OTX2
TNR
HTRA1
ST3GAL1
PCSK5
TSEN54
KCNK1
HMX1
COL9A1
TFAP2B
LINC00461
GBAS
TMEM161B
PHGDH
GCSH
PSPH
ZNF891
ZNF652
WDR5
WBP4
WAC
VKORC1L1
UMAD1
UBIAD1
UBE2L3
TRMT61B
TRAPPC3
TRAF3IP1
TLL1
TES
TANGO2
SYT13
SYNJ2
SUPT3H
STAG1
ST7L
SPTSSA
SPTBN1
SPCS3
SMIM12
SMG6
SLC25A22
SIX3
SH3PXD2A
SEL1L
SCFD2
SCAMP1
RPLP2
RPA3
RORA
RELN
RALGPS1
RALB
PTCD2
PSMC3
PRRT1
PPP2R3A
POU6F2
PNPLA2
PKN2
PKIA
PCLO
PCBP3
OXR1
OXA1L
ODF2L
NUP160
NUDT7
NSF
NR1H3
NDUFAF6
MVB12B
MTOR
MOCS2
MGMT
MFF
METTL15
ME3
MCPH1
Fovea
Peripheral
Rods
Cones
BP
HC
AC
RGC
Muller
Astro
MicG
Endo
MAPK8IP3
LSM4
LRIF1
KREMEN1
KBTBD7
ISCU
ICA1
IBTK
HS3ST1
HIVEP3
HDLBP
H1F0
GTF2B
FYN
FKBPL
FKBP9
FEZ2
FERMT2
FER
FAM213A
EXOSC10
EGLN3
EFHC1
DLC1
CWC25
CRK
COL6A1
CDO1
CAPZA1
CAMTA2
C4orf32
C11orf49
BRE
BRD3
BRAP
BCAS3
AURKAIP1
ATP13A2
ATG12
AFF3
ABCA6
MT−ND4
NFE2L2
PDE4DIP
RPL24
C1GALT1C1
TRIP12
SMARCA2
SRD5A3
KIF21A
TUBB3
SPG7
POLG
NDUFS1
MTPAP
CISD2
AUH
IFT43
TTC28
GRN
KCTD7
CLN8
CLN6
CLN5
TPP1
PPT1
IKBKG
WDR34
TMEM231
CEP41
PDE6D
PAX6
OPTN
GPR143
TMEM67
PEX6
PEX5
PEX19
PEX16
PEX14
PEX10
IFT80
GNB1
ZFP82
UTRN
UCHL3
UBXN2A
TBC1D5
TBC1D4
STAT2
SLK
SLC16A7
SEC11C
RBFOX1
POP4
OSCP1
NRXN3
NNT
MYT1L
MYSM1
MRPS15
MRPL19
MPRIP
INSR
INPP4B
HS6ST3
GPR158
GPATCH2
FAXC
COMMD6
COMMD1
CEP162
AKT3
AGAP1
ZNF280D
VPS13C
TXNRD2
TSC22D2
TMEM181
TMCO1
SLC6A15
SIX6
PRKAG2
PNPT1
NEAT1
MGAT5
MEIS1
MADD
LSM8
LRP12
LPP
LMO4
GNB1L
GMDS
FBXO32
FARS2
FAM120B
EXOC2
DST
CADM2
CACNA2D1
CACNA1A
BNIP1
ATXN2
ARHGEF12
ANKH
UQCRFS1
TSN
TRAPPC9
TOMM40
TMEM97
TLE4
TDRP
SYN3
SV2C
STOX2
Expression
Strength
2
1
0
−1

### Figure S3-2

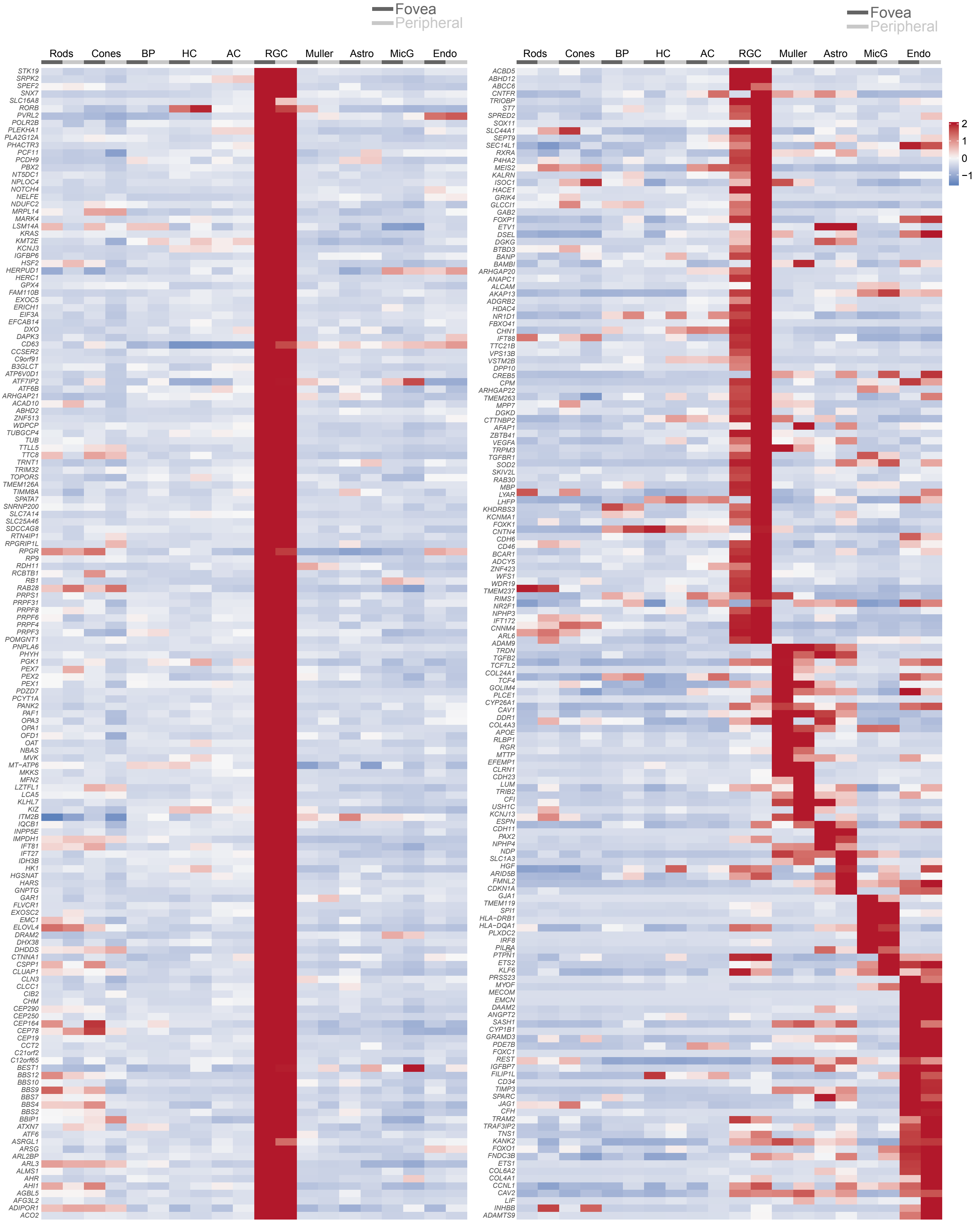
Fovea
Peripheral
Rods
Cones
BP
HC
AC
RGC
Muller
Astro
MicG
Endo
STK19
SRPK2
SPEF2
SNX7
SLC16A8
RORB
PVRL2
POLR2B
PLEKHA1
PLA2G12A
PHACTR3
PCF11
PCDH9
PBX2
NT5DC1
NPLOC4
NOTCH4
NELFE
NDUFC2
MRPL14
MARK4
LSM14A
KRAS
KMT2E
KCNJ3
IGFBP6
HSF2
HERPUD1
HERC1
GPX4
FAM110B
EXOC5
ERICH1
EIF3A
EFCAB14
DXO
DAPK3
CD63
CCSER2
C9orf91
B3GLCT
ATP6V0D1
ATF7IP2
ATF6B
ARHGAP21
ACAD10
ABHD2
ZNF513
WDPCP
TUBGCP4
TUB
TTLL5
TTC8
TRNT1
TRIM32
TOPORS
TMEM126A
TIMM8A
SPATA7
SNRNP200
SLC7A14
SLC25A46
SDCCAG8
RTN4IP1
RPGRIP1L
RPGR
RP9
RDH11
RCBTB1
RB1
RAB28
PRPS1
PRPF31
PRPF8
PRPF6
PRPF4
PRPF3
POMGNT1
PNPLA6
PHYH
PGK1
PEX7
PEX2
PEX1
PDZD7
PCYT1A
PANK2
PAF1
OPA3
OPA1
OFD1
OAT
NBAS
MVK
MT−ATP6
MKKS
MFN2
LZTFL1
LCA5
KLHL7
KIZ
ITM2B
IQCB1
INPP5E
IMPDH1
IFT81
IFT27
IDH3B
HK1
HGSNAT
HARS
GNPTG
GAR1
FLVCR1
EXOSC2
EMC1
ELOVL4
DRAM2
DHX38
DHDDS
CTNNA1
CSPP1
CLUAP1
CLN3
CLCC1
CIB2
CHM
CEP290
CEP250
CEP164
CEP78
CEP19
CCT2
C21orf2
C12orf65
BEST1
BBS12
BBS10
BBS9
BBS7
BBS4
BBS2
BBIP1
ATXN7
ATF6
ASRGL1
ARSG
ARL2BP
ARL3
ALMS1
AHR
AHI1
AGBL5
AFG3L2
ADIPOR1
ACO2
Fovea
Peripheral
Rods
Cones
BP
HC
AC
RGC
Muller
Astro
MicG
Endo
ACBD5
ABHD12
ABCC6
CNTFR
TRIOBP
ST7
SPRED2
SOX11
SLC44A1
SEPT9
SEC14L1
RXRA
P4HA2
MEIS2
KALRN
ISOC1
HACE1
GRIK4
GLCCI1
GAB2
FOXP1
ETV1
DSEL
DGKG
BTBD3
BANP
BAMBI
ARHGAP20
ANAPC1
ALCAM
AKAP13
ADGRB2
HDAC4
NR1D1
FBXO41
CHN1
IFT88
TTC21B
VPS13B
VSTM2B
DPP10
CREB5
CPM
ARHGAP22
TMEM263
MPP7
DGKD
CTTNBP2
AFAP1
ZBTB41
VEGFA
TRPM3
TGFBR1
SOD2
SKIV2L
RAB30
MBP
LYAR
LHFP
KHDRBS3
KCNMA1
FOXK1
CNTN4
CDH6
CD46
BCAR1
ADCY5
ZNF423
WFS1
WDR19
TMEM237
RIMS1
NR2F1
NPHP3
IFT172
CNNM4
ARL6
ADAM9
TRDN
TGFB2
TCF7L2
COL24A1
TCF4
GOLIM4
PLCE1
CYP26A1
CAV1
DDR1
COL4A3
APOE
RLBP1
RGR
MTTP
EFEMP1
CLRN1
CDH23
LUM
TRIB2
CFI
USH1C
KCNJ13
ESPN
CDH11
PAX2
NPHP4
NDP
SLC1A3
HGF
ARID5B
FMNL2
CDKN1A
GJA1
TMEM119
SPI1
HLA−DRB1
HLA−DQA1
PLXDC2
IRF8
PILRA
PTPN1
ETS2
KLF6
PRSS23
MYOF
MECOM
EMCN
DAAM2
ANGPT2
SASH1
CYP1B1
GRAMD3
PDE7B
FOXC1
REST
IGFBP7
FILIP1L
CD34
TIMP3
SPARC
JAG1
CFH
TRAM2
TRAF3IP2
TNS1
KANK2
FOXO1
FNDC3B
ETS1
COL6A2
COL4A1
CCNL1
CAV2
LIF
INHBB
ADAMTS9
2
1
0
−1

**Table S1**

| Donor ID | Retina ID | Donor Age | Donor sex | COD | Duration to process after death (hour) | 10X Kit |
|---|---|---|---|---|---|---|
| H1 | H1 | 74 | Male | Lung Cancer | 6 | V2 |
| H2 | H2R1 | 78 | Male | Metastatic Melanoma to brain | 14 | V2 |
| H2 | H2R2 | 78 | Male | Metastatic Melanoma to brain | 14 | V2 |
| H3 | H3 | 60 | Male | Left tonsillar squamous cell carcinoma metastatic to brain and left orbit | 6.5 | V2 |
| H4 | H4 | 64 | Male | Diffuse B cell lymphoma spread to thorax and epigastrum | 5 | V2 |
| H5 | H5 | 53 | Female | Interstitial Lung Disease | 5 | V2 |
| H9 | H9 | 76 | Male | Metastatic Melanoma | 6.5 | V3 |
| H11 | H11 | 65 | Male | Metastatic Melanoma | 3 | V3 |

**Table S2**

| Reference | Platform | Age | #Donors | # cells | Separate fovea/ macula and periphery | # clusters | Identify types within classes | Cells or nuclei |
|---|---|---|---|---|---|---|---|---|
| [12]Peng et al., 2019 | 10X, V2 | Adult | 1 | 2,383 | No | 9 | Yes | cells |
| [56]Hu et al., 2019 | Modified STRT | Fetal week 5-24 weeks | 19 embryos | 2,421 | No | 21 | No | cells |
| [55]Lukowski et al.,2019 | 10X, V2 | Adult | 3 | 20,009 | No | 17 | Yes | cells |
| [59]Voigt et al., 2019 | 10X, V3 | Adult | 3 | 8,217 | Yes | 17 | Yes | cryopreserved cells |
| [58]Menon et al., 2019 | 10X, V3 and Seq-Well | Adult | 6 | 23,432 | Yes | 9 | No | cells |
| [57]Liang et al., 2019 | ICELL8 | Adult | 3 | 5,873 | Yes | 7 | No | nuclei |
| [60]Sridhar et al., 2020 | 10X,V1,V2,V3 | Fetal | 4 embryos | 61,164 | Yes | 10 | No | Cells |
| This study | 10X, V2 and V3 | Adult | 8 | 85,000 | Yes | 58 | Yes | cells |
